# PNGaseA-mediated N-glycan stripping from peptides by infant-derived *Bifidobacterium bifidum*

**DOI:** 10.1101/2025.07.17.665090

**Authors:** Cassie R. Bakshani, Taiwo O. Ojuri, Simon G. Caulton, Keith Coughlan, Francesca Bottacini, Andrew L. Lovering, Douwe van Sinderen, Lucy I. Crouch

## Abstract

N-glycans are highly common sources of nutrition for human colonic-dwelling bacteria. These microbes have evolved a several methods to remove N-glycans from proteins; herein we describe the biochemical and structural characterisation of one such enzyme, a PNGaseA superfamily member produced by the infant-associated *Bifidobacterium bifidum* LMG13195. This PNGase was demonstrated to elicit activity against a wide variety of N-glycan structures yet exhibited a high preference for N-glycans attached to a peptide rather than to a denatured or native protein. This unusual specificity highlights how bacterial species tune their enzymology to different types of substrates. The structural characterisation of this PNGase reveals how its structure determines this specificity while being the first structure presented from the PNGaseA superfamily, revealing a unique ten-stand β-sheet cradling a canonical PNGase catalytic module.

## Introduction

N-glycans are posttranslational modifications on proteins, particularly common on proteins secreted extracellularly, such as antibodies^1^. In the context of the human colonic microbiota, the prevalence of this N-glycosylation means that they are ubiquitously available for these microbes to use as a nutrient source. N-glycans are both host-produced and diet-derived (from both animal and plant sources). Furthermore, fungal yeasts produce large mannan N-glycan structures as a part of their cell wall, so these will be available to the colonic microbes derived from yeast-fermented foods and from mycobiota species also residing in the colon^2^.

There are now several studies characterising the enzymology of N-glycan degradation by human-associated microbes: *Bacteroides thetaiotomicron* and mammalian complex N-glycans^3^, *Bacteroides thetaiotomicron* and yeast mannan^2^, *Streptococcus pneumoniae* and both high-mannose and complex N-glycans^4,5^, *Phocaeicola massiliensis* and plant/insect N-glycans^6^, and *Bifidobacterium* species and high-mannose N-glycans^7^. There are three families of enzymes that remove the N-glycan from the peptide component and one of these is the Peptide:N-glycosidases (PNGases), which hydrolyse between the protein and the core GlcNAc^1^. The two other families are glycoside hydrolase (GH) families 18 and 85, whose members hydrolyse between the two core GlcNAc sugars to separate the glycan from the protein^1^.

The infant colonic microbiota is dominated by species of *Bifidobacterium*, especially in breast-fed infants, at least partly due to their ability to directly or indirectly metabolise human milk oligosaccharides as their main carbohydrate and energy source^8^. However, N-glycans are also present in human milk and some species of *Bifidobacterium* have characterised GH18 and GH85 enzymes that are active against N-glycoproteins^9^. The presence of these enzymes in infant-associated species of *Bifidobacterium* indicate that N-glycans are also a source of nutrition. One notable observation showed growth of *B. longum subsp. infantis* ATCC 15697 on a mixture of complex N-glycans released from whey, including both sialylated and non-sialylated structures, of which the majority were degraded during growth^10^. In terms of high-mannose N-glycan breakdown, the full enzymatic machinery from a *Bifidobacterium longum* strain has been biochemically characterised^11^. Finally, N-glycan fragments have been observed in the faeces of breastfed infants and the dominant bacteria were species of *Bifidobacterium*^12^.

The ability of *Bifidobacterium* species to breakdown N-glycans as a nutrient source is thus evident from available literature, although the extent and importance of this activity has not yet been fully elucidated. To further enhance understanding in this area, we searched for possible PNGase enzymes encoded by genomes of infant-associated *Bifidobacterium* species and found a single putative candidate in two *B. bifidum* strains, one of which is *B. bifidum* LMG13195 (PNGaseA^Bif^). This enzyme belongs to the PNGaseA superfamily, and we used biochemical and structural characterisation to define specificity.

## Materials and Methods

### Bioinformatics

Initially, annotated PNGase enzymes were collated using Interpro, Uniprot and IMG databases^13–15^. This provided two candidates from *Bifidobacterium bifidum* LMG13195 and *Bifidobacterium leontopitheci*. Due to the presence of a PNGase being relatively uncommon to species of *Bifidobacterium*, the genetic context was then investigated. To analyse the variable region containing the gene encoding PNGaseA^Bif^ (*pngaseA^bif^*) across multiple *B. bifidum* genomes, the genbank files for 18 complete *Bifidobacterium bifidum* strains were downloaded from the National Centre for Biotechnology Information (NCBI) website. The initial search was filtered to include only complete genomes, resulting in the selection of the 18 genomes included in the analysis (accessed on 4th July 2025). The downloaded genbank files were uploaded to artemis (v16.0.5) and gene clusters were manually curated by searching for the actetate kinase and phosphoketalase genes, which flank the desired cluster. Once located, gene clusters were saved as a genbank file and uploaded to cagecat clinker1 for analysis^16^. The output graph was then refined to show only the unique clusters based on visual similarity and the number of strains found to contain each cluster was recorded on the figure. A detailed summary of all genomes analysed can be found in Supplementary Table 3. Alphafold3 was used to model interactions between PNGaseA^Bif^ and different peptides^17^.

Putative signal sequences were identified using SignalP 6.0^18^. The identities between different protein sequences were determined using Clustal Omega using full sequences^19^. The CAZy database was used to assess the GH families present in different *B. bifidum* strains^20^.

### Cloning and recombinant expression

The gene, *pngaseA^bif^*, was amplified from genomic DNA and cloned into pET28b (Novagen) using restriction sites NdeI and XhoI. The recombinant plasmid was transformed into TUNER cells (Novagen) and plated onto Luria-Bertani (LB) broth containing 50 μg/mL kanamycin and incubated overnight at 37 °C. Cells were then transferred into 1 L of LB medium (in a 2 L flask) and incubated at 37 °C until mid-exponential phase with shaking at 180 rpm. Cultures were then cooled to 16 °C, isopropyl β-D-thiogalactopyranoside (IPTG) was added to a final concentration of 0.2 mM, and the cultures were incubated overnight at 150 rpm and 16 °C in an orbital shaking incubator. These cells were harvested and recombinant His-tagged proteins were purified from the cell-free extracts using immobilised metal affinity chromatography (IMAC) using a buffer containing 20 mM Tris and 300 mM NaCl (pH 8.0). The purity and size of the proteins was assessed by sodium dodecyl sulfate-polyacrylamide gel electrophoresis (SDS-PAGE). The SDS-PAGE gels used were precast 4-16 % gradient (Bio-Rad) and stained with Coomassie Brilliant Blue (ReadyBlue® Protein Gel Stain, Sigma-Aldrich) to visualise total protein. The concentrations of recombinant enzymes were determined using absorbance at 280 nm using a Nanodrop and their calculated molar extinction coefficients. The other recombinant enzymes used in this report were expressed and purified using the same methods – BT3990 (*Bacteroides thetaiotaomicron*), BT0455 (*Bacteroides thetaiotaomicron*) and PNGaseL (*Flavobacterium akiainvivens*)^2,21,22^.

### Substrate preparation

Denatured forms of the substrates were prepared by boiling for 10 minutes in a heat block. Partially degraded substrates were prepared by first feeding the substrate to *Vibrio coralliilyticus* ATCC BAA-450, which only utilises the polypeptide, leaving the N-glycan attached to peptide fragments. *Vibrio coralliilyticus* ATCC BAA-450 was grown from glycerol stocks in marine broth (Difco) at 30 °C with shaking at 180 rpm. The following day, cells (1:100 dilution) were transferred to 5 mL of 1:1 2X marine M9 (M9 supplemented with 15 g/L NaCl) and 2X substrate (20 mg/mL) and grown at 30 °C with shaking at 180 rpm for 24 hours. Cultures were then centrifuged at 4,500 rpm for 5 minutes to pellet cells. The resultant supernatant was retained, filter sterilised and the fragmented glycoprotein was purified using HyperSep™ Hypercarb™ SPE Cartridges (Thermo Fisher Scientific).

### Enzyme assays

Screening for activity was carried out against bovine RNaseB (R7884, Sigma-Aldrich), bovine α1acid glycoprotein (G3643, Sigma-Aldrich) and bovine fetuin (F2379, Sigma-Aldrich) in native, denatured and partially degraded/digested forms. Horseradish peroxidase (77332, Sigma-Aldrich) in native and denatured forms was also used. Assays were carried out in 20 mM 4-morpholinepropanesulfonic acid (MOPS), pH 7.0, at 37 °C, with a final glycoprotein concentration of 10 mg/mL. The final enzyme concentration was 5 μM. The SDS-PAGE gels were precast 4-16 % gradient (Bio-Rad) and either stained with Coomassie Brilliant Blue (ReadyBlue® Protein Gel Stain, Sigma) to visualise total protein or Schiff’s Fuchsin (Pierce Glycoprotein Staining Kit) to visualise just glycoprotein.

### Thin-layer chromatography

9 μL of recombinant enzyme and whole cell assays were spotted onto silica plates. The plates were resolved in running buffer containing butanol/acetic acid/water in a 2:1:1 and 1:1:1 ratio for high-mannose N-glycan substrates and complex N-glycan substrates, respectively. They were resolved twice, with drying in between, and stained using a diphenylamine-aniline-phosphoric acid stain^23^.

### High-performance anion exchange chromatography-pulsed amperometric detection (HPAEC-PAD)

To analyse N-glycan release, samples were separated using a CarboPac PA300-4µm anion exchange column with a PA300G-4µm guard using a Dionex ICS-6000 (Thermo Fisher) and detected using PAD. The flow was 0.25 mL min^-1^ and the buffers were A: water, B: 200 mM NaOH, C: 50 mM NaOH and 25 mM NaOAc, and D: 100 mM NaOH and 250 mM NaOAc.

The elution conditions were 0-4 mins 80 % A, 19.5 % B and 2 % C, 4-20 mins had a linear increase of B and C to 20 and 60 %, respectively. This was followed by 20-50 mins where buffer D was increased linearly to 100 % and remained at that level from 50-60 mins to clean the column. Software used was the Chromeleon Chromatography Data System. All data were obtained by diluting assays 1/10 in sterile MilliQ H2O prior to injection.

### Crystallisation, data collection and structure determination

To prepare PNGaseA^Bif^ for crystallisation a second purification step after IMAC was applied. Size-exclusion chromatography on a Superdex 200 26/200 column (Cytivia) using an AKTA Pure FLPC system (GE Healthcare Life Sciences) was carried out with a buffer containing 25 mM HEPES, 200 mM NaCl (pH 8). SDS-PAGE gels were used to determine the pure fractions, which were then pooled and concentrated. PNGaseA^Bif^ was initially screened using commercial kits, using a sitting drop vapour diffusion method and set up using a Mosquito liquid handling robot (TTP Labtech). The crystallisation experiments were composed of 0.1 μL of reservoir solution and 0.2 μL of protein (25 mg/mL). A second drop of 0.1 μL reservoir solution was also included. The trays were sealed and incubated at 18 °C. Crystal clusters first formed in PACT condition 0.2 M NaI, 0.1 M Bis-tris propane, 20 % PEG3350 (w/v), pH 7.5. The clusters were fragmented via centrifugation with Microseed beads in the same reservoir solution that they formed. The seed stock was soaked in 10 mM chitobiose and was used to set up a fresh PACT screen. The crystal that diffracted successfully was formed in PACT condition 46 - 0.1 M Tris pH 8.0, 20 % PEG 6000 (w/v), 0.2 M Magnesium chloride. For cryoprotection, an additional 25 % PEG400 (w/v) was added to the crystallisation liquor prior to flash-freezing crystals in liquid nitrogen. Data from the crystals were collected at the European Synchrotron Radiation Facility (ESRF) using the ID30B beamline. Data were processed using autoPROC and STARANISO. The structure was solved by molecular replacement using coordinates of PNGaseA^Bif^ model generated using Alpha-fold colab^24^. The structures were phased and refined using Phenix^25^. Manual building was carried out in COOT^26^. The structure was analysed and figures generated using Pymol^27^. Consurf was used to assess conservation of residues over the PNGaseA^Bif^ structure^28^. Despite soaking with chitobiose, no ligand was present in the active site.

## Results

### Identifying PNGaseA^Bif^

To investigate N-glycan utilisation by species of human-associated *Bifidobacterium* species, Interpro, Uniprot and IMG databases were screened for putative PNGase-encoding sequences, and this resulted in the identification of a predicted PNGase gene present in the genome of *Bifidobacterium bifidum* LMG13195 (the enzyme is denoted here as PNGaseA^Bif^, while its corresponding gene is referred to here as *pngaseA^bif^*)^13–15^. Intriguingly, PNGaseA^Bif^ belongs to the PNGaseA superfamily, which is not currently associated with nutrient acquisition in bacterial species^13^.

In the InterPro database, the current putative PNGaseA superfamily members are present in 1404, 710 and 94, eukaryotic, bacterial and archaeal species, respectively^13^. Within the eukaryotes, 1009 species are fungi (mainly Ascomycota and Basidiomycota), 311 species are flowering plants, while the remainder currently represent unclassified species. For the bacteria, 571 of the species with a putative PNGaseA-encoding sequence are Actinobacteria, where 444 of these are species of *Streptomyces*. The rest of the bacterial species represent a wide variety of microorganisms that are members of the Pseudomonadati kingdom. Of these bacterial sequences, there is only one additional putative PNGaseA enzyme (other than that specifying PNGaseA^Bif^) associated with a member of the *Bifidobacterium* genus, *Bifidobacterium leontopitheci* (A0A6I1GFR2) isolated from Golden-headed Lion Tamarin and Goeldi’s monkey^29^. However, this putative PNGaseA member elicits just 25 % protein sequence identity to PNGaseA^Bif^.

To investigate the genetic context of *pngaseA^bif^* in the genome of *Bifidobacterium bifidum* LMG13195 and to assess why this gene is apparently absent from other members of this species, we analysed corresponding chromosomal regions in several *B. bifidum* strains from the NCBI genome database (Supplementary Table 1). Eighteen complete genomes were available at the time of this comparative analysis and revealed that *pngaseA^bif^*is part of a highly variable region flanked by two conserved regions (Supplementary Figures 1 and 2). In *B. bifidum* LMG13195, this variable region contains the PNGaseA^Bif^ gene and various transposase-encoding genes, suggesting that *pngaseA^bif^* was acquired by horizontal gene transfer. Overall, eight different types of inserts were detected in this variable region for these eighteen genomes (Supplementary Figure 2). Through this analysis, a second *B. bifidum* strain (Bg41221_3D10), isolated from a human infant in Bangladesh in a recent project, has a very similar gene pattern in this variable region, including a *pngA* of high identity to that found in LMG13195^30^ (Supplementary Figure 2).

Initial analysis of the PNGaseA^Bif^ amino acid sequence revealed several characteristics (Figure 1). Sequence alignments of PNGaseA^Bif^ to PNGaseF superfamily representatives highlighted the likely eight-stranded β-sandwiches in the first N-terminal half of the PNGaseA^Bif^ sequence (Supplementary Figure 1). There is a large region after this which had no homology to the PNGaseF representatives, labelled ‘X’ and then a proline/serine/glycine/glutamine-rich region that is likely a linker. There is then a LPXTG-like motif, which may be a site for sortase-dependent anchoring to the cell wall, and finally a predicted transmembrane region at the C-terminus^31^. The signal sequence is predicted to be a Sec signal peptide (SPI), indicating that PNGaseA^Bif^ is produced by *Bifidobacterium bifidum* LMG13195 as an extracellular, membrane-anchored enzyme^18^.

**Figure 1.**
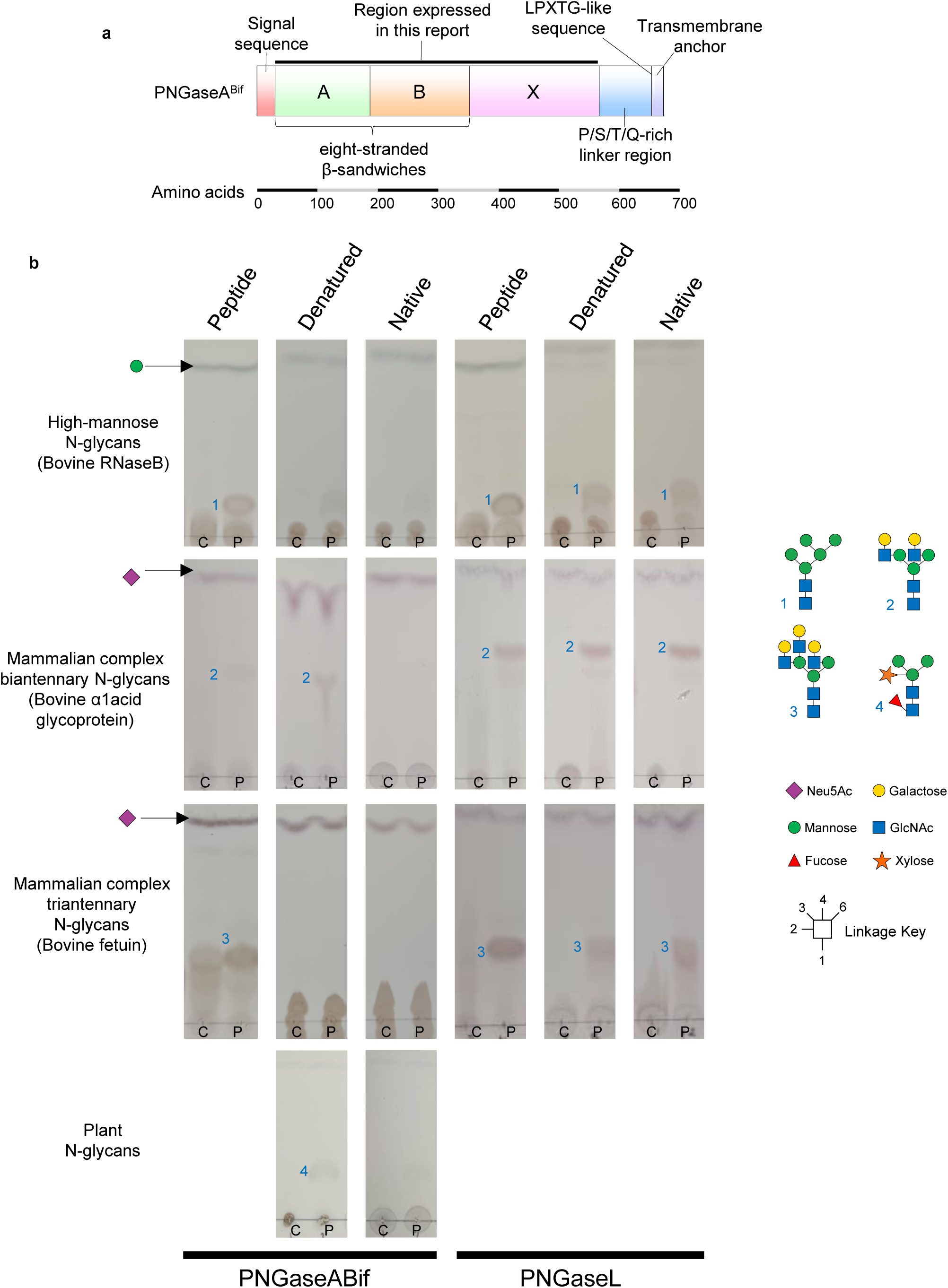
PNGaseA^Bif^ domain structure and activity. **a)** The two eight-stranded β-sandwiches predicted to comprise the catalytic module are labelled A and B. The section after this, labelled X, was not assigned any annotation by databases. Towards the C-terminus of the protein a region of low complexity was highlighted by searches and is comprised of predominantly proline, threonine, serine and glutamine, which suggests this area is a linker region. The C-terminus is predicted to be a transmembrane region, suggesting that the enzyme is anchored to the cell surface of *B. bifidum*. The black line represents the section cloned in this study and the signal sequence is predicted to be SPI Sec. b) Recombinantly-expressed PNGaseL and PNGaseA^Bif^ were tested against glycoprotein substrates with different types of N-glycans and with the protein in the native form, pre-treated with boiling (denatured) and with the protein mostly digested away (peptide). The release of N-glycans by the PNGases are visualised by thin layer chromatography. The results presented here are with the addition of α-mannosidase or α-sialidase where applicable, but full data sets are presented in Supplementary Figures X-X). C and P indicate the control and PNGase samples, respectively.

### Investigating the specificity of PNGaseA^Bif^

Previously characterised members of the PNGaseA superfamily exhibit a preference for N-glycans that are covalently linked to only a peptide^32^. Therefore, assays were designed to test specificity towards both the N-glycan and protein components of these substrates. Three glycoprotein substrates were initially screened that have three different types of N-glycans: high-mannose (bovine RNaseB), mammalian biantennary complex (bovine α1acid glycoprotein) and mammalian triantennary complex (bovine fetuin). The protein part of the substrate was also prepared in three different ways – no pre-treatment (native), boiling (denatured) and by removing most of the protein to leave just a fragment (peptide). To produce the N-glycosylated peptide fragments, the native N-glycosylated substrates were provided as a nutrient source to the highly proteolytic coral pathogen, *Vibrio coralliilyticus*^33^. This species uses most of the protein as a nutrient source and the remaining N-glycan on a short peptide is purified for assays. In addition to this, the high-mannose N-glycan and complex N-glycan substrates were also pretreated with an α1,2-mannosidase (BT3990) and a broad-acting α-sialidase (BT0455), respectively, to fully explore potential activity (Figure 1 and Supplementary Figures 3-5)^2,22^.

The results were initially visualised by thin-layer chromatography (TLC) and sodium dodecyl sulfate–polyacrylamide gel electrophoresis (SDS-PAGE), which monitor for released glycan and changes in glycoprotein size, respectively. SDS-PAGE gels were stained for protein and glycoprotein using Coomassie brilliant blue or Schiff’s Fuchsin, respectively (Figure 1 and Supplementary Figures 3-5). PNGaseL was included as a positive control and activity was observed under all conditions^21^. PNGaseA^Bif^ was determined to possess activity primarily towards N-glycans attached to just a peptide, some activity against the denatured substrates, and no activity against the native version. To analyse the released glycans from these assays in more detail, high-performance anion exchange chromatography with pulsed amperometric detection (HPAEC-PAD; Figures 2 and 3) was used. For the high-mannose N-glycoprotein, released glycans could be detected for denatured protein with both the positive control and PNGaseA^Bif^. Notably, high mannose N-glycans of a variety of sizes (Man5-8) were released by PNGaseA^Bif^. For the N-glycan on a peptide fragment, the high background means that the peaks corresponding to the different glycans are not clearly identifiable, however, the peak corresponding to Man5 has been highlighted. Furthermore, the chromatograms for the samples where PNGaseA^Bif^ was included are different to the control, whereas the PNGaseL chromatograms look akin to the control. The data suggests that the addition of an α1,2-mannosidase, under these assay conditions, does not affect the efficiency of the control or PNGaseA^Bif^. The HPAEC-PAD data from assays against glycoproteins with mammalian complex N-glycans also support the conclusions drawn from the TLCs. The activity observed for PNGaseA^Bif^ when the N-glycan is attached to a peptide fragment is considerably higher. The addition of α-sialidase, under these assay conditions, does not appear to affect the efficiency of the control or PNGaseA^Bif^.

**Figure 2.**
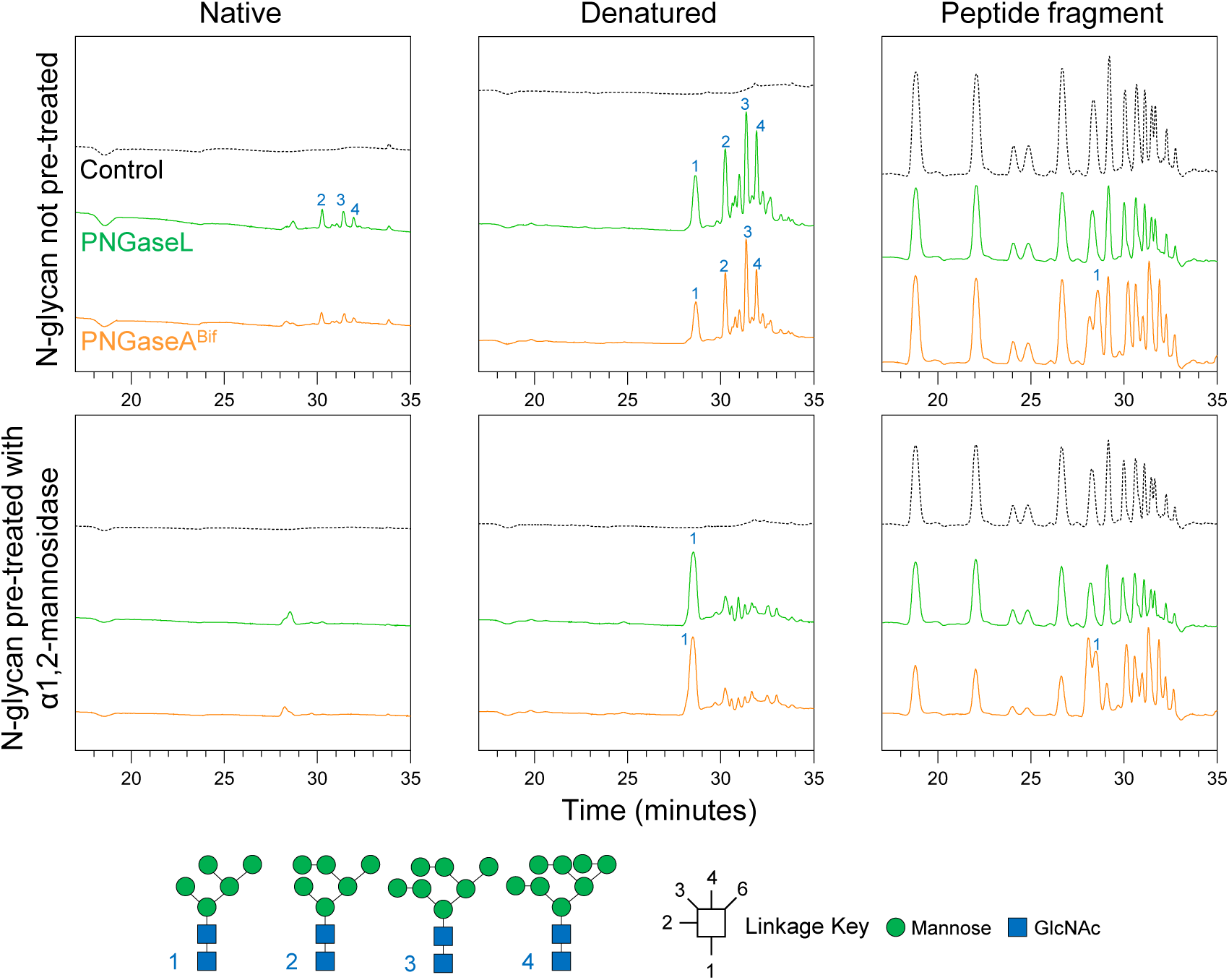
The activity of PNGaseA^Bif^ against high-mannose N-glycans. Released N-glycans were assessed using HPAEC-PAD data. The top three panels assess a variety of high-mannose N-glycans structures and the bottom three panels assess when all α1,2-mannose sugars have been removed to leave just Man5.

**Figure 3.**
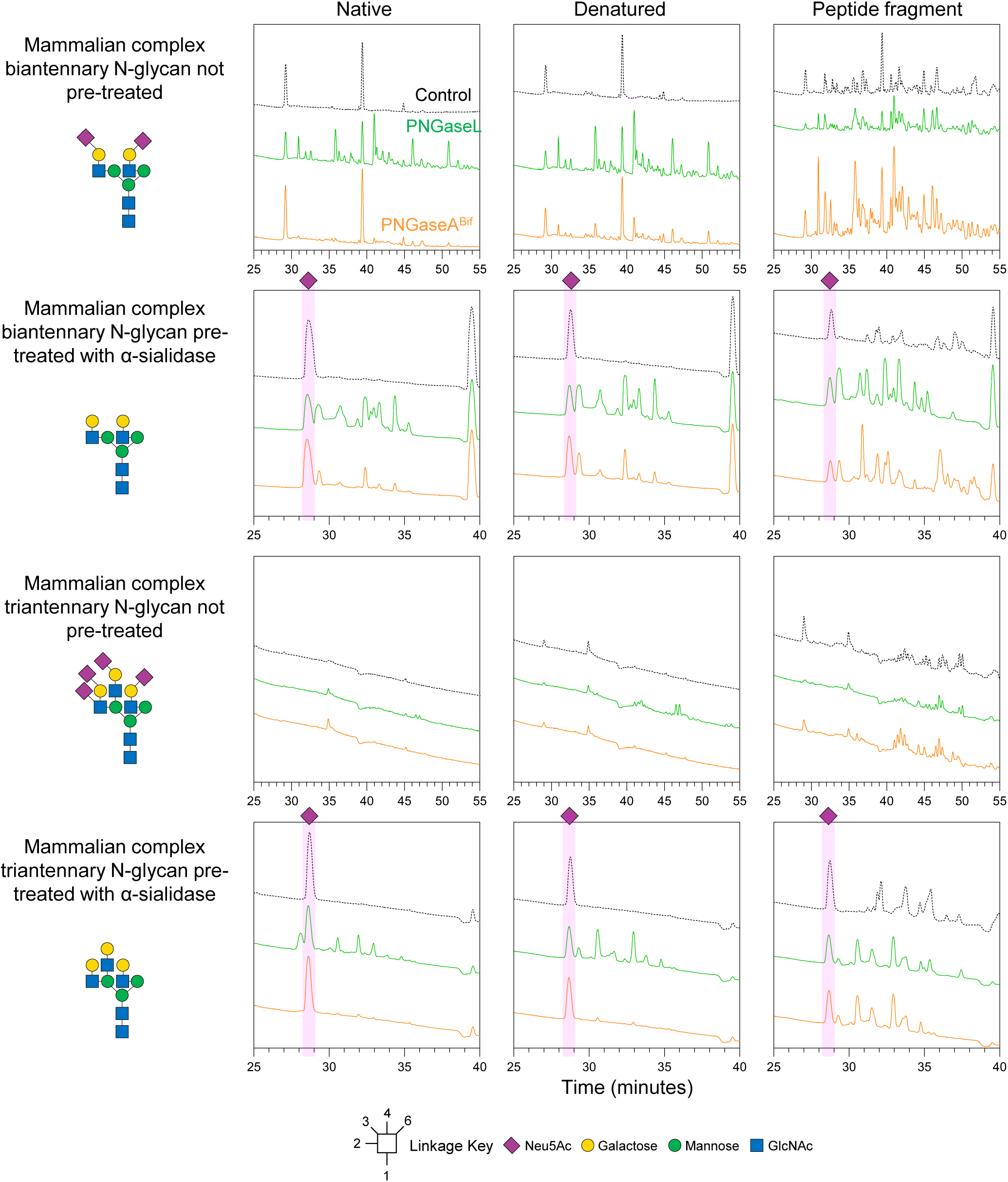
The activity of PNGaseA^Bif^ against mammalian complex N-glycans. Released N-glycans were assessed using HPAEC-PAD data. Substrates decorated with either biantennary or triantennary complex N-glycans were assessed and the addition of a broad acting α-sialidase was also used to explore specificity in more detail.

To assess the influence of α-fucosylation of the core GlcNAc on activity, we used the plant-derived glycoprotein, horseradish peroxidase, as a substrate. It was only possible to carry out these experiments under native and denatured conditions due to the nature of the proteins (not enough to test bacterial growth and subsequent purification). PNGaseA^Bif^ was able to release glycans from this substrate, suggesting that core α1,3-fucose can be accommodated in the active site. Activity against plant and insect type N-glycans has been observed previously for both PNGaseA and PNGaseF superfamily members^1^.

### Crystal structure of the PNGaseA^Bif^ from *B. bifidum*

To investigate the structural basis for the unusual specificity displayed by PNGaseA^Bif^ we determined the crystal structure to 1.9 Å (Figure 4, Supplementary Figures 6-9, Supplementary Table 2). The PNGaseA^Bif^ catalytic module comprises two eight-stranded β-sandwich domains and this overlays well with the solved structures from the PNGaseF superfamily. To demonstrate this, superimposition of the crystal structure of archetypal PNGaseF from *Elizabethkingia meningoseptica* (1PNF) and PNGaseA^Bif^ shows an RMSD of 4.8 over 248 residues (Figure 4). Strikingly, however, in the PNGaseA^Bif^ structure there is an additional large ten-strand β-sheet closely associated with this core catalytic module which cradles the entirety of one side of the catalytic module (Figure 4). This is absent in all other solved PNGase structures^6,34,35^.

**Figure 4.**
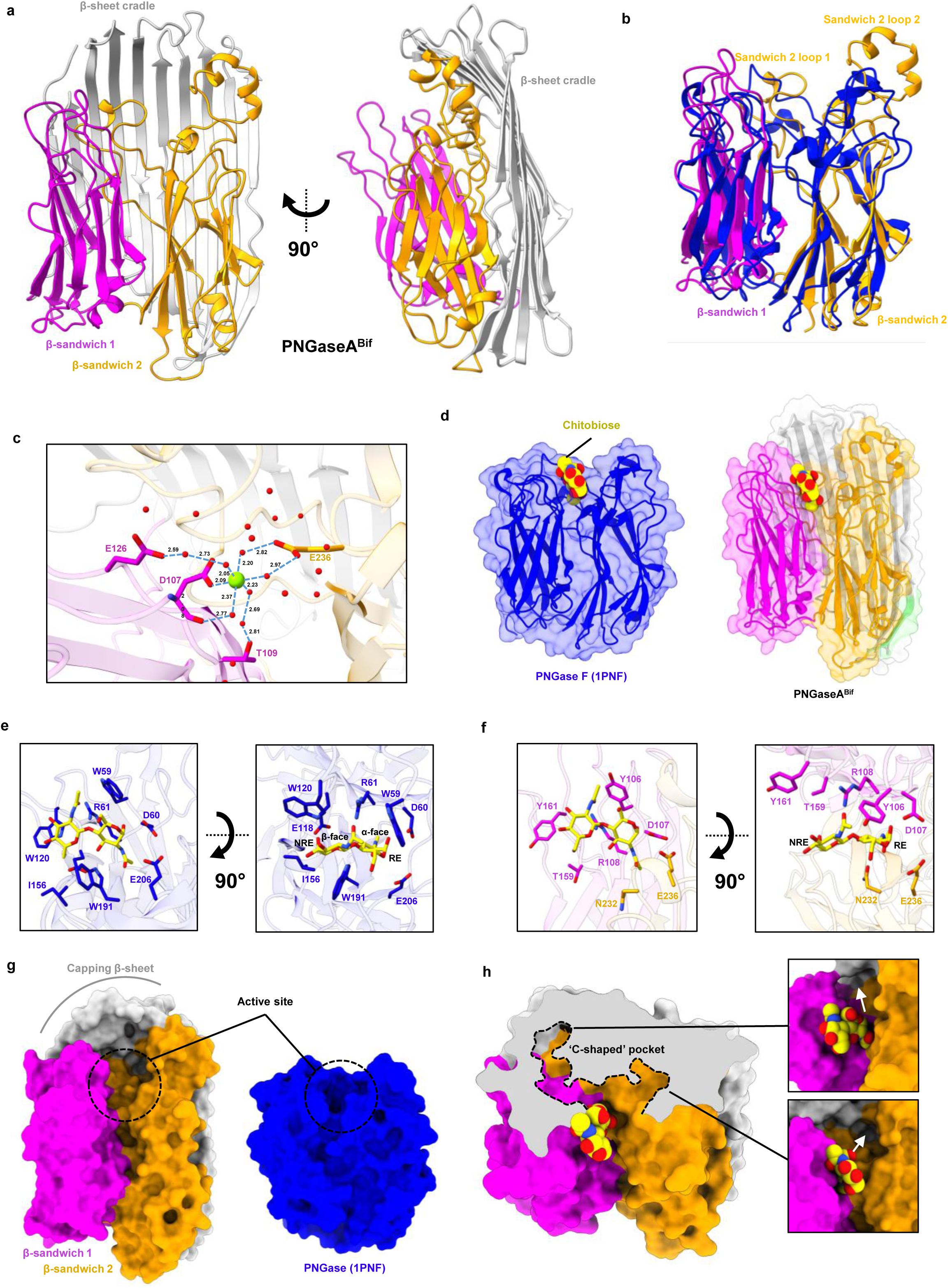
Structure of PNGaseA^Bif^ characterised in this study. **a)** A cartoon representation of the PNGaseA^Bif^ structure shown from two different angles with the β-sandwiches coloured in magenta and gold and the β-sheet cradle in grey. b) An overlay of the eight-stranded β-sandwich catalytic domains from PNGaseA^Bif^ and PNGaseF (1PNF, blue). c) Residues and water molecules within the active site of PNGaseA^Bif^ are shown. The Mg^2+^ ion (green) is coordinated by the carboxylate oxygen of the side chain and 5 waters in octahedral geometry. The 5 waters are then hydrogen bonded directly or indirectly to the side chains of glutamic acid 126, glutamic acid 236, threonine 109, and the backbone carbonyl of aspartic acid 107. Dashed lines are shown for direct and indirect interactions, with distances shown in Angstroms. d) Surface representations of PNGaseF (blue) and PNGaseA^Bif^ (pink/gold/silver) overlaid on top of a cartoon representation. Chitobiose from 1PNF is in yellow highlights the position of the active site in both enzymes. e) The active site of PNGaseF 1PNF with chitobiose (yellow) shown from two different angles. f) The active site of PNGaseA^Bif^ with the chitobiose (yellow) from 1PNF overlaid. g) Surface representations of PNGaseA^Bif^ and PNGaseF to emphasise the different substrate-binding regions. h) A cross-section of the active site to show the C-shape pocket that likely accommodates short peptides (dotted line) and two closeup views of the active site, looking into the two pockets where the short peptides will likely be accommodated.

Notably, a magnesium ion was observed in the active site of this structure, and it is coordinated by waters molecules and four residues, both directly and indirectly, in an octahedral geometry (D107, T109, E126, E236; Figure 4 and Supplementary Figure 6). The equivalent residues to D107 and E236 in PNGaseF (D60 and E206) have previously been shown to be essential for catalytic activity^35^. To our knowledge, metal has not been implicated in the catalytic activity of PNGases and the importance of magnesium for activity was not explored further here. Similar metal binding has not been observed in other structural data for PNGase enzymes from other organisms. The structure of PNGaseF (1PNF) was solved with chitobiose in the active site and the superimposition of PNGaseF and PNGaseA^Bif^ shows that this magnesium ion is in very close proximity to the reducing end of the core GlcNAc (Supplementary Figure 6).

The overlay of the PNGaseF 1PNF structure and PNGaseA^Bif^ also allows comparison of the residues in the active sites. Interactions with the α-face of the core GlcNAc and the β-face of the second GlcNAc show significant similarities for four out of five residues (Figure 4). Both have two aromatic residues that are tryptophan (W59 & W120) and tyrosine (Y106 & Y161) in PNGaseF and PNGaseA^Bif^, respectively. Both structures also have an arginine and aspartate on this side of the disaccharide that overlay well. The residue that differs between the two structures is a glutamic acid (E118) and a threonine (T159) in PNGaseF and PNGaseA^Bif^, respectively, nearest to C6 of the second GlcNAc. In PNGaseF, there is evidence that E118 blocks N-glycans with α1,3-fucose core decorations so PNGaseF cannot cleave them^34^. In comparison, T159 in the PNGaseA^Bif^ structure is relatively smaller and contributes to a more open pocket around the C6 and C3 of the non-reducing end and reducing end GlcNAc sugars, respectively. This supports the biochemical data obtained for PNGaseA^Bif^ where release of N-glycans from horseradish peroxidase was observed and these glycans predominantly have α1,3-fucose core decorations^6^ (Figure 1). Opposite the conserved aspartic acid (D107), there is a conserved glutamic acid (E236) and, as stated above, both have been shown to be essential for catalytic activity^35^. Finally, on the other side of the disaccharide (β-face of the core GlcNAc and the α-face of the second GlcNAc) this area is much more open for PNGaseA^Bif^ compared to PNGaseF. PNGaseF has an isoleucine and tryptophan pinning the disaccharide into the active site, whereas an asparagine is in the place of this tryptophan in PNGaseA^Bif^ with no equivalent residue to the isoleucine.

Analysis of the area around the active site provides insights into why PNGaseA^Bif^ prefers N-glycans on a peptide. Either side of where the reducing end of the core GlcNAc would be positioned are two long pockets that together form a “C” shape (Figure 4). This is likely where a short peptide would be accommodated during catalysis. In comparison to this, the PNGaseF structure is very open around the catalytic site, likely allowing it to accommodate N-glycans attached to larger protein components. PNGaseA^Bif^ B-factors suggest that the loops connecting the β-sheets of the cradle near the active site are dynamic, so the structure may open and close to accommodate differently sized substrates (Supplementary Figure 7). This would explain why PNGaseA^Bif^ can also remove N-glycans from some denatured glycoproteins.

An electrostatic overlay was applied to the PNGaseA^Bif^ structure to visualise the charge of the surface (Supplementary Figure 7). Interestingly, one of the pockets is highly negatively charged and the entrance to the other is highly positively charged. In theory, this would support the accommodation of the charged amino- and carboxy-termini of a peptide and support the overall proposal that the function of the C-shaped pocket is to accommodate a peptide. To test the potential for this pocket to accommodate peptide, Alphafold3 was used to model in potential peptides (Supplementary Figure 8). The results indicate that the negatively charged end of the pocket (formed by sandwich 2 loop 2) accommodates the N-terminus of the peptide and a maximum of three or four residues at the N-terminus would allow the correct positioning of an asparagine in the active site. Furthermore, the C-terminal part of the peptide was accommodated in the positively charged part of the C-shaped pocket and threaded through to the open-end of this pocket. The C-terminal part of the peptide is clasped tightly by the β-sheet cradle and the loops of β-sandwich 1 (Supplementary Figure 8).

To assess the conservation of the protein sequences across the structure, we used ConSurf (Supplementary Figure 9). The results show that the highest conservation of residues is in and around the active site and in two areas on either side of the active site where the β-sheet cradle meets the active site module. On one side there are two cysteines that do not form a disulphide bond in this structure (Supplementary Figure 10) while on the other side there are two highly conserved arginine residues. These two arginine amino acids are also the residues generating the positive charge at the entrance to one of the pockets.

Notably, the C-shaped double pocket of PNGaseA^Bif^ is formed where the catalytic module and the β-sheet cradle meet (Figure 4 and Supplementary Figure 8), therefore the presence of this sheet in other putative structures will likely indicate a preference for N-glycans attached to short peptides. To investigate this, we assessed models of other PNGaseA across species from Fungi, Bacteria, Archaea, Oomycota and Plant Kingdoms (Supplementary Figure 11). The models predict that the two eight-stranded β-sandwich structures of the catalytic module and the large β-sheet cradle are highly conserved features in all structures assessed. There are some minor differences, such as a large extra loop in the *Candida viswanathii* structure and the incomplete cradle in the *Cordyceps confragosa* structure. The putative PNGaseA from *Bifidobacterium leontopitheci* was the only example of a putative structure where a small additional module was observed.

### Carbohydrate-Active enzyme conservation across *B. bifidum* strains

The presence of a gene encoding a PNGaseA appears to be relatively uncommon among members of the *Bifidobacterium* genus from the searches carried out during this work. To understand the wider biological context of where PNGaseA^Bif^ fits into the glycan breakdown capacity of *B. bifidum* LMG13195, the conservation of other Carbohydrate-Active enZyme (CAZyme) families was assessed across different strains with different origins of isolation (Figure 5). The presence of putative GH family members, their frequency within a genome, and the conservation of different GH enzymes across different strains was assessed for host-associated CAZyme families (Table 2). Overall, the frequency of a given GH family is consistent across strains, especially for host-associated enzyme families. The most striking aspect of this analysis was the very high conservation of CAZymes between strains for the different CAZyme primary sequences (Supplementary Table 3). Those enzymes that were assessed had no less than 90 % identity.

**Figure 5.**
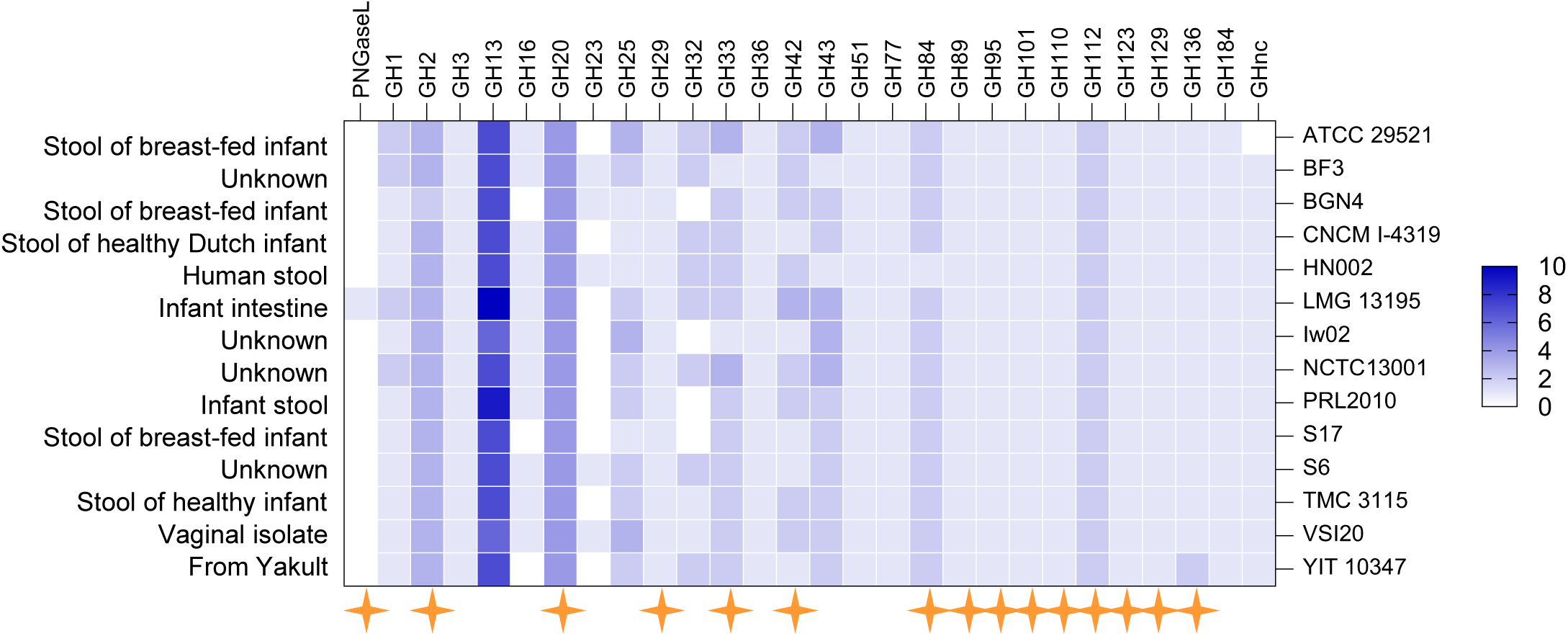
Frequency of putative CAZymes across different *Bifidobacterium bifidum* strains. The heat map shows how many different GH and PNGase families are present across different strains (right). The origin of the strain is indicated on the left and the orange four-pointed star indicates the enzyme families associated with host glycan breakdown.

The highest represented GH family is 13, with between six and ten putative enzymes. These enzymes likely degrade starch and other α-glucan polysaccharides, in addition to roles in energy storage within the cell. No GH13 enzymes from *B. bifidum* are currently characterised. The second most prevalent family is GH20 with a consistent four enzymes across strains and >90 % identity in protein sequence. This family often plays a critical role in host glycan degradation in human commensal microbes, and, for example, it is the largest represented GH family in the mucinophilic human colonic commensal *Akkermansia mucinophila*^36^. The GH20 family members typically have β-HexNAc’ase activities and those that have been characterised for *B. bifidum* show distinct specificities^37–40^. For the other host-associated GH family members, there is typically only a single highly conserved enzyme, for example GH29, GH89, GH95 and GH110.

### A model of host glycan breakdown by *B. bifidum* LMG13195

To construct a model detailing how *B. bifidum* LMG13195 breaks down different host glycans, we used a combination of cellular localisation prediction and current literature (Figure 6). *B. bifidum* strains have a large number of CAZymes predicted to be on their cell surface relative to other *Bifidobacterium* species, and these likely act to trim down relatively large host glycans to substrates that can be imported for further breakdown by CAZymes residing inside the cell. A range of both exo- and endo-acting CAZymes are predicted to be on the surface.

**Figure 6.**
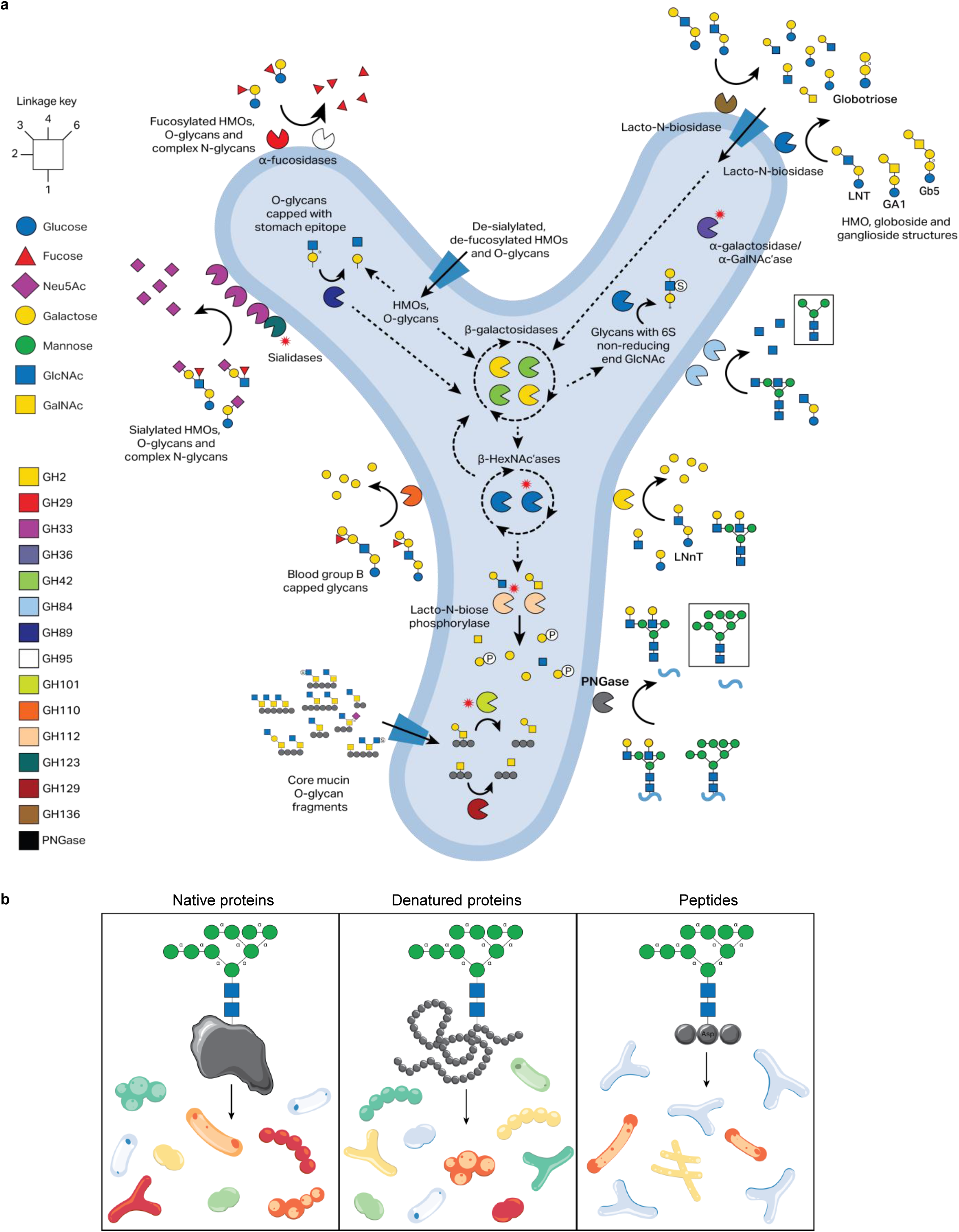
A model for the breakdown of host glycans by *Bifidobacterium bifidum* LMG13195. **a)** This model includes the breakdown of human milk oligosaccharide structures, O-glycans, N-glycans and complex glycosphingolipid structures. The information is predominantly taken from the literature and the red twelve-pointed stars indicate where biochemical characterisations are currently unknown. The enzymes are coloured according to their family and further information is available in Supplementary Table 3. **b)** A model for how the state of the protein component of a glycoprotein drives enzyme specificity and, therefore, directs nutrition to different microbial species. Microbial species expressing PNGaseA-type enzymes will likely be primarily be accessing N-glycans on peptides (right panel). Whereas N-glycans linked to denatured proteins (central panel) or native proteins (left panel) will likely be accessed by microbes expressing PNGaseF superfamily members or GH18/85 family members, respectively.

Regarding the exo-acting enzymes, there are three α-sialidases (GH33), two α-fucosidases (GH29 and GH95), an α-galactosidase (GH110), a β-galactosidase (GH2), two β-HexNAc’ases (GH84) and a putative α-GalNAc’ase (GH123) predicted to be on the surface. The published activities are collated in Supplementary Table 3 and from this information a hypothesis can be proposed about the order different host glycans are broken down into their component sugars. All of these enzymes, bar the GH123, have had some level of specificity defined, which means that a fairly accurate glycan utilisation model can be proposed for the CAZymology taking place on the surface of *B. bifidum*. The first two CAZymes to act will be the GH33 and GH110 enzymes that will remove α-linked sialic acid and blood group B α-galactose^41–44^, respectively. Sialic acid decorates many different host glycans and blood group B is present in 0-30 % of the population, depending on genetics. The two GH29 and GH95 enzymes have been shown to act on α1,3/4- and α1,2-linkages, respectively, and human milk oligosaccharides, N-glycans and O-glycans are all commonly decorated with α-fucose^45,46^. Notably, there are no α-sialidases or α-fucosidases predicted to be inside the cell, so this process is compartmentalised to outside the cell, which means that the import machinery likely plays a selective role here. The GH2 β-galactosidase has been characterised as having β1,4 specificity and demonstrated to act on Lacto-N-*neo*tetraose, lactose and N-glycans^47^. The two GH84 family members have both been shown to have broad β-GlcNAc’ase activity, including against N-glycans^39^. Finally, GH123 family members are currently all characterised as having β-GalNAc activity, which is most common in ganglioside and globoside structures, but can be present in O- and N-glycans also.

The endo-acting CAZymes predicted to be on the surface of the cell include two Lacto-N-biosidases from families GH20 and GH136. The GH20 can remove β1,3 disaccharides from Lacto-N-tetraose, ganglioside GA1 and globoside Gb5^37^. The products are likely then imported for further degradation, although the globotriose product from the Gb5 may be further broken down by the GH110. The GH136 has been shown to have the ability to remove both Lacto-N-biose and LacNAc from Lacto-N-tetrose and Lacto-N-*neo*tetraose, respectively^48^. These are common human milk oligosaccharide and O-glycan structures and are likely imported, although products from both GH20 and GH136 enzymes with a β1,4-galactose at the non-reducing end may be hydrolysed by the GH2 β-galactosidase. Finally, in the case of *B. bifidum* LMG13195, the PNGaseA is predicted to also be on the surface and target N-glycans on short peptides which can then be sequentially broken down by the GH2 and GH84 enzymes. *B. bifidum* genomes do not encode GH families with α-mannosidase activities, so it is hypothesised that the mannose parts of these glycans are not accessed as a nutrient source but are instead cross-fed to other members of the microbiota.

Once the host glycans are imported into the cell, it is hypothesised that they can be broken down using the four β-galactosidase (2xGH2 and 2xGH42) and three β-GlcNAc’ases (GH20)^38–40,47^. Notably, one of these GH20 enzymes has specificity for 6S-GlcNAc common to mucins. In parallel with these enzymes, there are two GH112 putative Lacto-N-biose phosphorylases currently uncharacterised. These two GH112 enzymes have ∼ 85 % identity between them and both are highly conserved over the strains assessed.

Mucin O-glycans produced in the stomach are commonly capped with α1,4-GlcNAc at the non-reducing end and the GH89 from *B. bifidum* has been characterised to have specificity towards this sugar^49^. The reason why this enzyme is intracellular in comparison the sialidases and fucosidases, for example, remains poorly understood. In terms of mucin breakdown, *B. bifidum* has a GH101 and GH129 predicted to be intracellular. These enzyme families are characterised to act on the T and Tn antigen of O-glycan cores, which implies that *B. bifidum* can import O-glycoproteins. Finally, a GH36 is also predicted to be inside the cell and this family has been shown to have α-galactosidase/GalNAc’ase activities. It would be logical that this enzyme is an α-GalNAc’ase, used to break down blood group A structures, but specificity is still to be defined^36^.

## Discussion

The colonisation of the infant colon by species of *Bifidobacterium* plays a vital and holistic role in the development of a healthy gut microbiota. A full mechanistic understanding of how these microbes colonise so successfully, particularly in the presence of breast milk, has yet to be fully elucidated. One of the main factors driving successful colonisation are the human milk oligosaccharides found in breast milk, but other factors, including N-glycans, also influence this process^50^.

In this study, we characterise a PNGaseA superfamily member from *B. bifidum* LMG13195 and determined its preference for N-glycans attached to peptides, rather than denatured or native protein. Conversely, PNGaseA^Bif^ could accommodate a broad selection of different N-glycan structures, including high-mannose, mammalian complex, and simple plant N-glycans with core α1,3-fucose. This highlights how the protein component of glycoprotein substrates, rather than the glycan component, can influence the types of substrates microbes can access, which has not been explored before in microbial nutrient acquisition (Figure 6). In the case of colonic-dwelling *B. bifidum*, the breakdown of the protein component of a glycoprotein would occur either by proteolytic microbes or the host digestive enzymes and the remaining N-glycan:peptide would be a substrate for a PNGaseA.

Furthermore, we solved the structure of PNGaseA^Bif^ to understand the origin of the observed specificity. This is the first structure of a PNGaseA superfamily member and shows some remarkable features linking structure to substrate niche. PNGaseA^Bif^ shows a unique β-sheet cradle holding a canonical catalytic module and this structural component contributes significantly to the specificity of PNGaseA^Bif^. From our analysis, we can propose that a C-shaped pocket accommodates peptides with N-glycans and modelling using Alphafold3 provides approximate parameters for the types and sizes of peptides that PNGaseA^Bif^ can accommodate.

Finally, a glycan utilisation model for *Bifidobacterium bifidum* LMG13195 was constructed and provides unique insight into the capability of this species to breakdown a wide variety of host-derived substrates. This model underlines what is known about the glycobiology of this species and what remains to be understood.

## Data availability

The atomic coordinates and structure factors have been deposited in the Protein Data Bank (ID code 9RZW).

## Acknowledgements

The work was funded by The Academy of Medical Sciences (SBF0061175), the Wellcome Trust and Royal Society Sir Henry Dale fellowship (224240/Z/21/Z) awarded to L.I.C. T.O.O. is funded by the BBSRC Midlands Integrative Biosciences Training Partnership (MIBTP) with his studentship in collaboration with industrial partners Ludger (Oxford, UK) awarded to L.I.C. DvS is supported by Research Ireland (formerly Science Foundation Ireland) Grant numbers SFI/12/RC/2273-P1 and SFI/12/RC/2273-P2. We would like to thank the European Synchrotron Radiation Facility for beamtime and staff on Beamline ID30b (19/4/2023).

---

**Supplementary Table 1.** *B. bifidum* strain information used to analyse the variable region where *pngaseA^bif^* resides in LMG13195.

| Accession no. | Strain Name | NCBI Submitter | Genbank | Refseq | Date Published | Date Accessed |
| --- | --- | --- | --- | --- | --- | --- |
| NZ_CP058603.1 | CNCM I-4319 | University College Cork | GCA_017894325.1 | GCF_017894325.1 | 09-Apr-2021 | 22-May-2025 |
| NZ_LR134344.1 | NCTC13001 | SC | GCA_900637095.1 | GCF_900637095.1 | 20-Dec-2018 | 22-May-2025 |
| NZ_AP012323.1 | JCM 1255 | Graduate School of Frontier Sciences, University of Tokyo | GCA_001025135.1 | GCF_001025135.1 | 24-Mar-2015 | 22-May-2025 |
| NZ_CP169560.1 | Bg41221_3D10 | Pathology and Immunology, Washington University School of Medicine in St. Louis | GCA_050308165.1 | GCF_050308165.1 | 14-May-2025 | 04-Jun-2025 |
| NZ_CP022723.1 | S6 | Korea Food Research Institute | GCA_003390735.1 | GCF_003390735.1 | 15-Aug-2018 | 22-May-2025 |
| NZ_LR698991.1 | MGYG-HGUT-02396 | EMG | GCA_902386775.1 | GCF_902386775.1 | 10-Aug-2019 | 04-Jun-2025 |
| NZ_CP149430.1 | lw02 | Peking University School and Hospital of Stomatology, Peking University | GCA_045290295.1 | GCF_045290295.1 | 26-Nov-2024 | 04-Jun-2025 |
| NZ_CP018757.1 | PRI 1 | Institute for Basic Science (IBS) and Pohang University of Science and Technology (POSTECH) | GCA_002845845.1 | GCF_002845845.1 | 20-Dec-2017 | 04-Jun-2025 |
| NZ_CP118071.1 | VSI20 | Seattle University | GCA_029011515.1 | GCF_029011515.1 | 06-Mar-2023 | 04-Jun-2025 |
| NZ_CP069279.1 | HN002 | Korea Research Institute of Bioscience and Biotechnology | GCA_016838705.1 | GCF_016838705.1 | 09-Feb-2021 | 04-Jun-2025 |
| NZ_AP031420.1 | i03-0019-A5 | Munehiro Furuichi Keio University school of medicine, Department of Microbiology and Immunology; Shinanomachi 35, Shinjuku-ku, Tokyo 160-8582, Japan | GCA_042848835.1 | GCF_042848835.1 | 13-Oct-2024 | 04-Jun-2025 |
| NZ_CP010412.1 | BF3 | Yonsei University | GCA_001281345.1 | GCF_001281345.1 | 14-Sep-2015 | 04-Jun-2025 |
| NZ_AP018132.1 | TMC 3115 | Graduate School of Biostudies,<br>Kyoto University | GCA_003573895.1 | GCF_003573895.1 | 03-Oct-2017 | 04-Jun-2025 |
| NZ_AP024712.1 | YIT 10347 | Yakult Central Institute | GCA_020892075.1 | GCF_020892075.1 | 03-Nov-2021 | 04-Jun-2025 |
| NC_014638.1 | PRL2010 | Laboratory of Probiogenomics,<br>Department of genetics,<br>university of Parma, Italy | GCA_000165905.1 | GCF_000165905.1 | 02-Nov-2010 | 04-Jun-2025 |
| NC_017999.1 | BGN4 | Korea Research Institute of<br>Bioscience and Biotechnology<br>(KRIBB) | GCA_000265095.1 | GCF_000265095.1 | 05-Jun-2012 | 04-Jun-2025 |
| NC_014616.1 | S17 | Institute of Microbiology and<br>Biotechnology, University of Ulm | GCA_000164965.1 | GCF_000164965.1 | 19-Oct-2010 | 04-Jun-2025 |
| AP018131.1 | JCM 7004 | Graduate School of Biostudies,<br>Kyoto University | GCA_003573955.1 | - | 14-Sep-2017 | 22-May-2025 |

**Supplementary Table 2.** Data collection and refinement statistics.

|  |  |
| --- | --- |
|  | PNGase |
| Wavelength |  |
| Resolution range | 68.46 - 1.9 (1.93 - 1.9) |
| Space group | C 1 2 1 |
| Unit cell | 161.981 81.037 124.642 90 127.83 90 |
| Total reflections | 228381 |
| Unique reflections | 85024 (3341) |
| Multiplicity | 2.7 (2.2) |
| Completeness (%) | 85.6 (62.9) |
| Mean I/sigma(I) | 6.4 (1.4) |
| Wilson B-factor | 24.96 |
| R-merge | 0.113 (0.562) |
| R-meas | 0.140 (0.712) |
| R-pim | 0.081 (0.429) |
| CC1/2 | 0.985 (0.667) |
| Non-patterson peak | - |
| Reflections used in refinement | 84900 (1652) |
| Reflections used for R-free | 4118 (64) |
| R-work | 0.1863 (0.3326) |
| R-free | 0.2209 (0.4248) |
| Number of non-hydrogen atoms | 9446 |
| macromolecules | 8274 |
| ligands | 58 |
| solvent | 1114 |
| Protein residues | 1078 |
| RMS(bonds) | 0.013 |
| RMS(angles) | 1.35 |
| Ramachandran favored (%) | 96.83 |
| Ramachandran allowed (%) | 2.89 |
| Ramachandran outliers (%) | 0.28 |
| Rotamer outliers (%) | 1.19 |
| Clashscore | 7.14 |
| Average B-factor | 32.16 |
| macromolecules | 30.84 |
| ligands | 55.42 |
| solvent | 40.75 |

Statistics for the highest-resolution shell are shown in parentheses.

**Supplementary Table 3.** A summary of the biochemical characterisations of CAZymes from *Bifidobacterium bifidum* LMG13195.

| Glycoside hydrolase family | CBMs in the same ORF | Locus tags | GenBank number | Predicted localisation (ref Sig6) | Published activity? | Known or putative substrate | Conservation across *B. bifidum* strains |
| --- | --- | --- | --- | --- | --- | --- | --- |
| GH1 |  | BBJK_01169 | BBA47815.1 | No signal sequence | No published activity |  |  |
| GH1 |  | BBJK_01541 | BBA48062.1 | No signal sequence | No published activity |  |  |
| GH2 | | BBJK_01655 | BBA48139.1 | No signal sequence | Lactose not GOS<sup>47</sup> | β-galactose | >95 % identical across strains |
| GH2 | | BBJK_00662 | BBA47484.1 | No signal sequence | Broad activity<sup>47</sup> | β-galactose | >97 % identical across strains |
| GH2 | CBM32 | BBJK_02214 | BBA48495.1 | SPI Sec | Lactose, LacNAc, LNnT<sup>47</sup> | β-galactose | >90 % identical across strains |
| GH3 |  | BBJK_02595 | BBA48753.1 | No signal sequence | No published activity |  |  |
| GH13 | | BBJK_00049 | BBA47087.1 | | No published activity | α-linked glucose | |
| GH13 | | BBJK_00050 | BBA47088.1 | | No published activity | α-linked glucose | |
| GH13 | | BBJK_00052 | BBA47089.1 | | No published activity | α-linked glucose | |
| GH13 | | BBJK_01640 | BBA48130.1 | | No published activity | α-linked glucose | |
| GH13 | | BBJK_01063 | BBA47754.1 | | No published activity | α-linked glucose | |
| GH13 | | BBJK_00541 | BBA47407.1 | | No published activity | α-linked glucose | |
| GH13 | | BBJK_03218 | BBA49152.1 | | No published activity | α-linked glucose | |
| GH13 | CBM48 | BBJK_00828 | BBA47596.1 | | No published activity | α-linked glucose | |
| GH13 | CBM48 | BBJK_02658 | BBA48796.1 | | No published activity | α-linked glucose | |
| GH13 | CBM48 | BBJK_02688 | BBA48814.1 | | No published activity | α-linked glucose | |
| GH16 | fragment | BBJK_02086 | BBA48422.1 | No signal sequence | No published activity |  |  |
| GH20 | | BBJK_00630 | BBA47460.1 | SPI Sec | LnbB lacto-N-biosidase, LNT, GA1, Gb5<sup>37</sup> | β-HexNAc | >92 % identical across strains |
| GH20 | 2x CBM32 | BBJK_00776 | BBA47560.1 | No signal sequence | Bbhl β1,3lacto-N-triose,N-glycan GlcNAcβ1,2Man<sup>38,39</sup> | β-HexNAc | >99 % identical across strains |
| GH20 | CBM32 | BBJK_03032 | BBA49032.1 | No signal sequence | BbhII 6S-GlcNAc'ase<sup>40</sup> | β-HexNAc | >99 % identical across strains |
| GH20 | | BBJK_01451 | BBA48002.1 | No signal sequence | No published activity | β-HexNAc | >98 % identical across strains |
| GH25 | 3xCBM13 | BBJK_01493 | BBA48030.1 |  | No published activity | Peptidoglycan |  |
| GH25 | CBM50 | BBJK_02952 | BBA48982.1 |  | No published activity | Peptidoglycan |  |
| GH29 | | BBJK_00483 | BBA47373.1 | SPI Sec | α1,3/4 fucosidase<sup>45</sup> | α-fucose | >96 % identical across strains |
| GH32 |  | BBJK_02265 | BBA48527.1 |  | No published activity |  |  |
| GH32 |  | BBJK_02267 | BBA48528.1 |  | No published activity |  |  |
| GH33 | | BBJK_01327 | BBA47922.1 | SPI Sec | SiaBb2. Broad activity<sup>42-44</sup> | α-sialic acid | |
| GH33 | | BBJK_01326 | BBA47921.1 | SPI Sec | SiaBb1. Also has esterase activity<sup>51</sup> | α-sialic acid | |
| GH36 |  | BBJK_01363 | BBA47949.1 | No signal sequence |  |  | >98 % identical across strains |
| GH42 | | BBJK_00477 | BBA47368.1 | | B216_08266, Galβ1,3/4/6 linkages, 3FL<sup>47</sup> | β-galactose | >99 % identical across strains |
| GH42 | | BBJK_01679 | BBA48156.1 | | B216_08730, Galβ1,4 linkages<sup>47</sup> | β-galactose | >99 % identical across strains |
| GH43 |  | BBJK_01432 | BBA47992.1 |  | No published activity |  |  |
| GH43 |  | BBJK_02021 | BBA48381.1 |  | No published activity |  |  |
| GH43 |  | BBJK_02023 | BBA48382.1 |  | No published activity |  |  |
| GH51 |  | BBJK_01448 | BBA48000.1 |  | No published activity |  |  |
| GH77 | | BBJK_00833 | BBA47600.1 | | No published activity | α-linked glucose | |
| GH84 | 3x<br>CBM32 | BBJK_00753 | BBA47544.1 | SPI Sec | BbhIV, Core 2, β1,3lacto-N-triose, N-glycan GlcNAcβ1,2Man<sup>39</sup> | β-GlcNAc | >98 % identical across strains |
| GH84 | | BBJK_00930 | BBA47665.1 | SPI Sec | BbhV, Core 2, β1,3lacto-N-triose, N-glycan GlcNAcβ1,2Man<sup>39</sup> | β-GlcNAc | >98 % identical across strains |
| GH89 | 4x<br>CBM32 | BBJK_00728 | BBA47530.1 | No signal sequence | Stomach group<sup>49</sup> | α-GlcNAc | >98 % identical across strains |
| GH95 | | BBJK_01721 | BBA48185.1 | SPI Sec | α1,2 fucosidase<sup>46</sup> | α-fucose | >97 % identical across strains |
| GH101 | CBM32 | BBJK_01844 | BBA48266.1 | No signal sequence | No published activity | Mucin cores | >98 % identical across strains |
| GH110 | CBM51 | BBJK_00385 | BBA47302.1 | SPI Sec | Acts on Blood group B<sup>41</sup> | α-galactose | >97 % identical across strains |
| GH112 |  | BBJK_01792 | BBA48232.1 | No signal sequence | No published activity | Lacto-N-biose | >99 % identical across strains |
| GH112 |  | BBJK_03100 | BBA49080.1 | No signal sequence | No published activity | Lacto-N-biose | >98 % identical across strains |
| GH123 | GH33 | BBJK_00867 | BBA47625.1 | SPI Sec | No published activity | β-GalNAc<br>α-sialic acid | >98 % identical across strains |
| GH129 | | BBJK_01138 | BBA47796.1 | No signal sequence | Acts on Tn antigen<sup>52</sup> | Exo-α-GalNAc'ase | >98 % identical across strains |
| GH136 |  | BBJK_02171 | BBA48469.1 | SPI Sec | LNT and LNnT<sup>48</sup> | Human milk oligosaccharides | >92 % identical across strains |
| GHnc |  | BBJK_02041 | BBA48392.1 | No signal sequence | No published activity |  |  |
| PNGaseA |  |  |  | SPI Sec | This work | N-glycans |  |

**Supplementary Figure 1.**
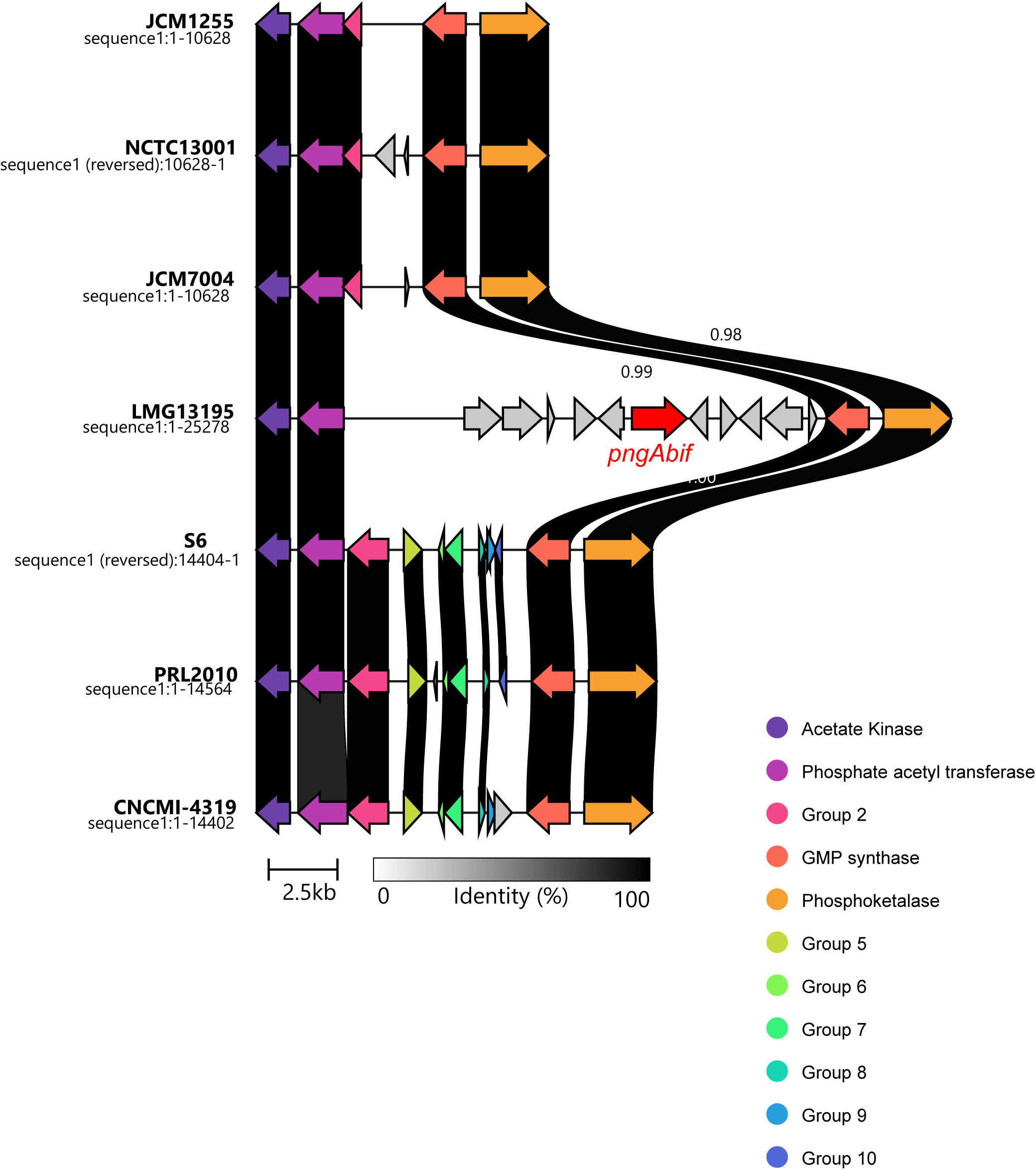
Genetic context of *pngAbif.* The gene cluster where *pngAbif* is found (red) in LMG13195 is shown in comparison to other *B. bifidum* strains. Out of the eighteen complete genomes, eight different proposed gene clusters could be identified. The conservation of the different genes is displayed through the greyscale mapping between the genes, which indicates near 100 % identity between all genes assesses. The flanking genes for these gene clusters consist of acetate kinase and a phosphate acetyl transferase (purples), typically on the left flank, as well as a GMP synthase and phosphoketalase (oranges), typically on the right flank. Inversions of this gene cluster are possible, and one such inverted cluster can be observed in the PRI 1 gene cluster, which was re-orientated for purpose of illustration here.

**Supplementary Figure 2.**
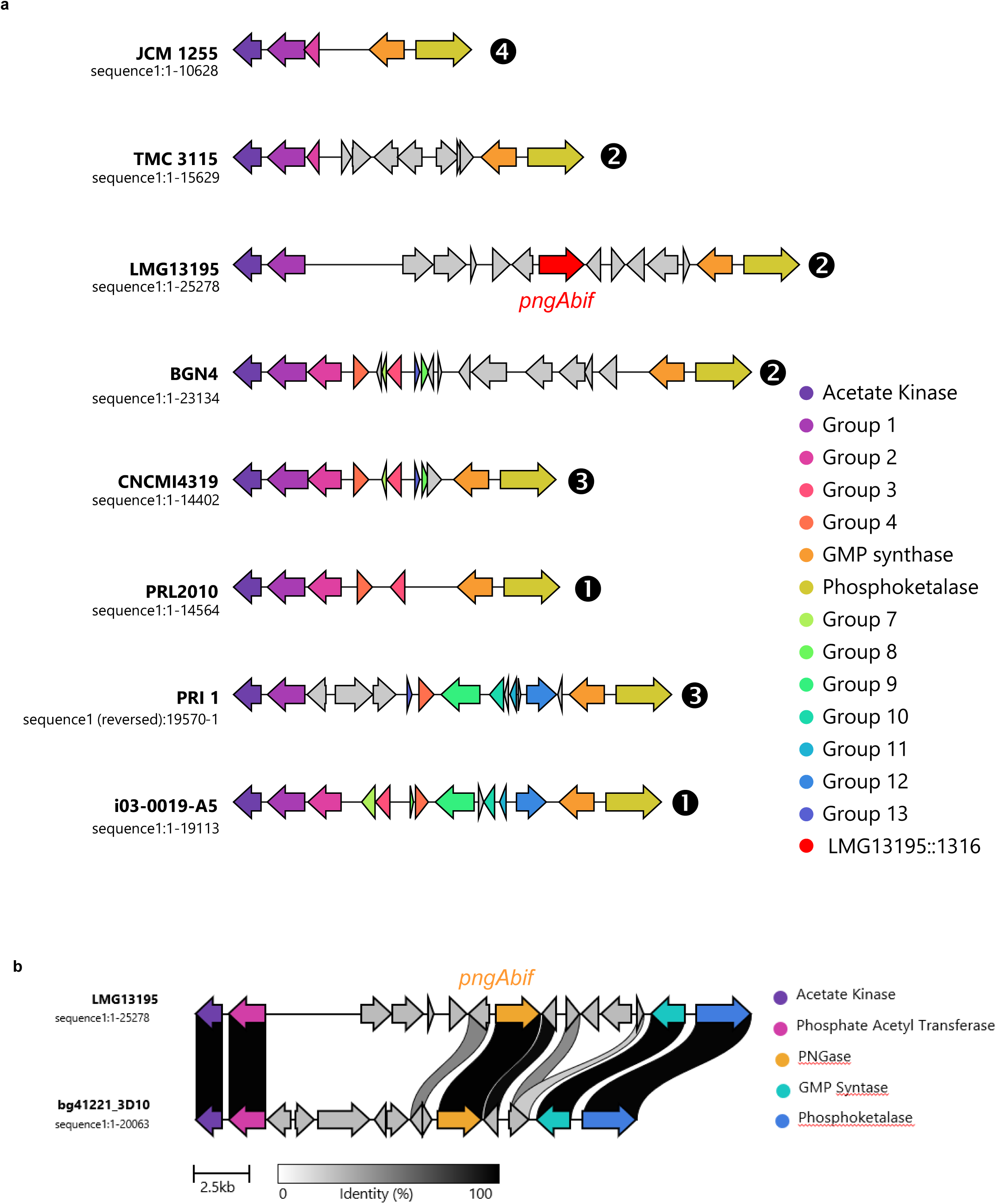
Variable region in *Bifidobacterium bifidum* genomes. **a)** Out of the eighteen complete genomes, eight different proposed gene clusters could be identified. The *pngAbif* is highlighted in red and homologues are indicated by matching colours. The number of times a particular genome structure was observed is indicated by the white numbers in black circles. b) A comparison of the two B. bifidum genomes encoding genes for putative PNGaseA enzymes. There is some variation between the two, but *pngAbif* is highly conserved.

**Supplementary Figure 3.**
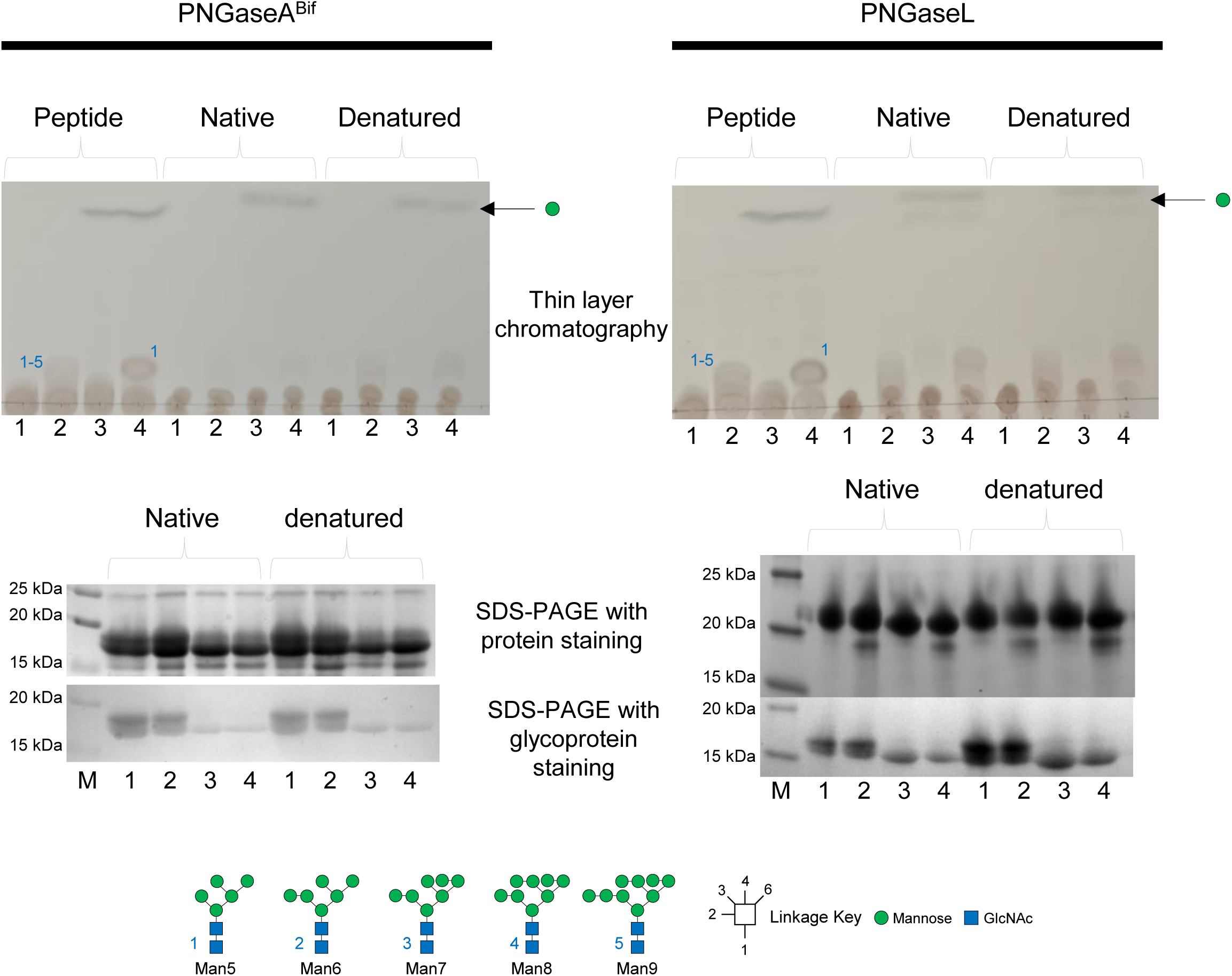
Biochemical Characterisation of PNGaseA^Bif^ against high-mannose N-glycans. Bovine RNaseB was used to assess the specificity of PNGaseA^Bif^ (left) for high-mannose N-glycans and PNGaseL was used as a positive control (right). The peptide was provided to the PNGase enzymes in three forms – native, denatured (boiling), and peptide (*Vibrio coralliilyticus* treated). The impact of removing the α1,2-linked mannose from the non-reducing ends of the glycans was also investigated using BT3990 from *Bacteroides thetaiotaomicron*. The N-glycan release was assessed through thin layer chromatography (top) and change in glycoprotein size was monitored through SDS-PAGE with staining for protein (middle) and staining for glycoprotein (bottom). 1: control, 2: PNGase, 3: α1,2-mannosidase, 4: PNGase and α1,2-mannosidase. M: marker. The *Vibrio coralliilyticus-*derived fragments are too small to resolve by SDS-PAGE.

**Supplementary Figure 4.**
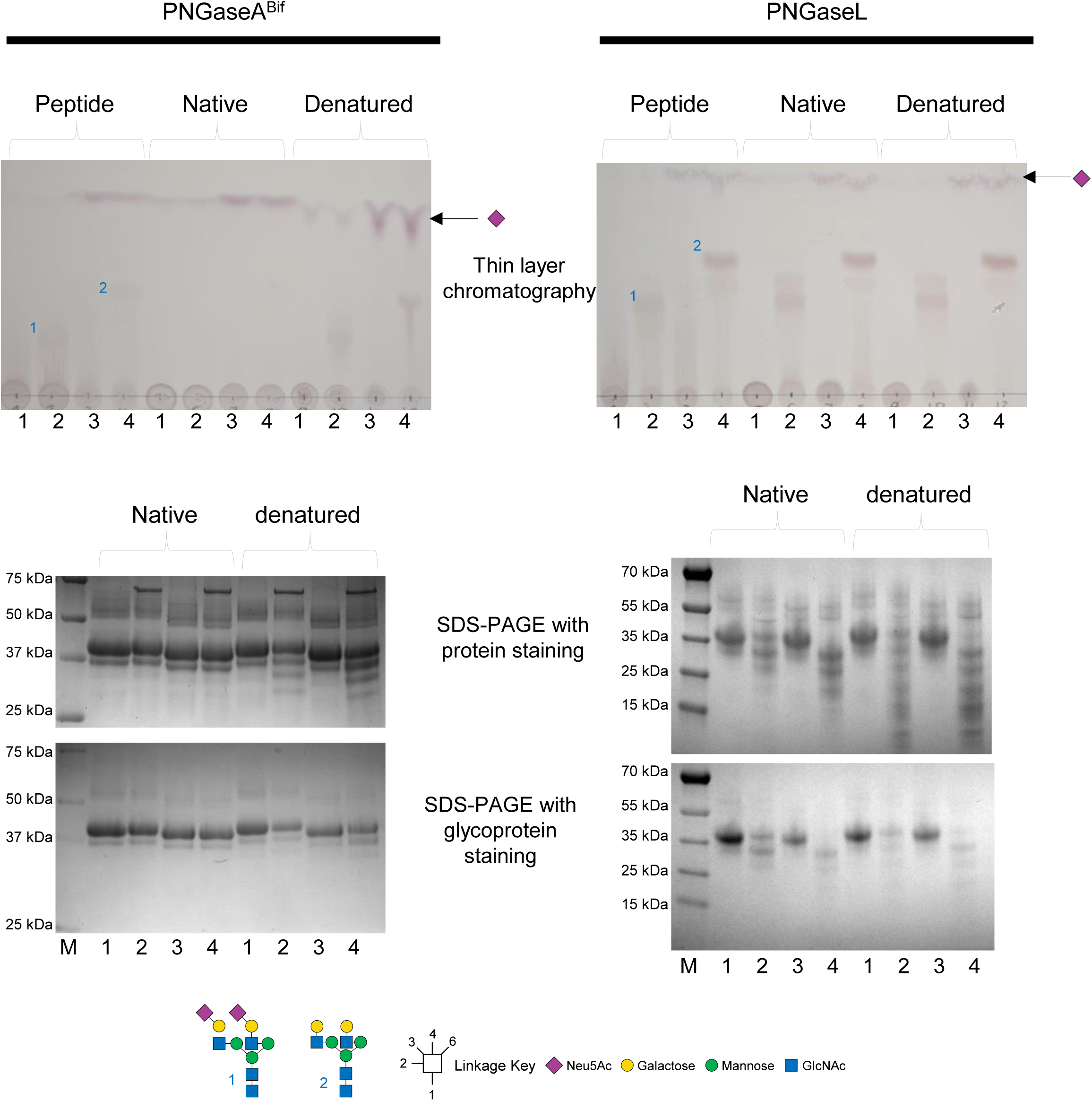
Biochemical Characterisation of PNGaseA^Bif^ against mammalian biantennary complex N-glycans. Bovine α1acid glycoprotein was used to assess the specificity of PNGaseA^Bif^ (left) for mammalian biantennary complex N-glycans and PNGaseL was used as a positive control (right). The peptide was provided to the PNGase enzymes in three forms – native, denatured (boiling), and peptide (*Vibrio coralliilyticus* treated). The impact of removing the α-linked sialic acid from the non-reducing ends of the glycans was also investigated using BT0455 from *Bacteroides thetaiotaomicron*. The N-glycan release was assessed through thin layer chromatography (top) and change in glycoprotein size was monitored through SDS-PAGE with staining for protein (middle) and staining for glycoprotein (bottom). 1: control, 2: PNGase, 3: α-sialidase, 4: PNGase and α-sialidase. M: marker. The *Vibrio coralliilyticus-*derived fragments are too small to resolve by SDS-PAGE.

**Supplementary Figure 5.**
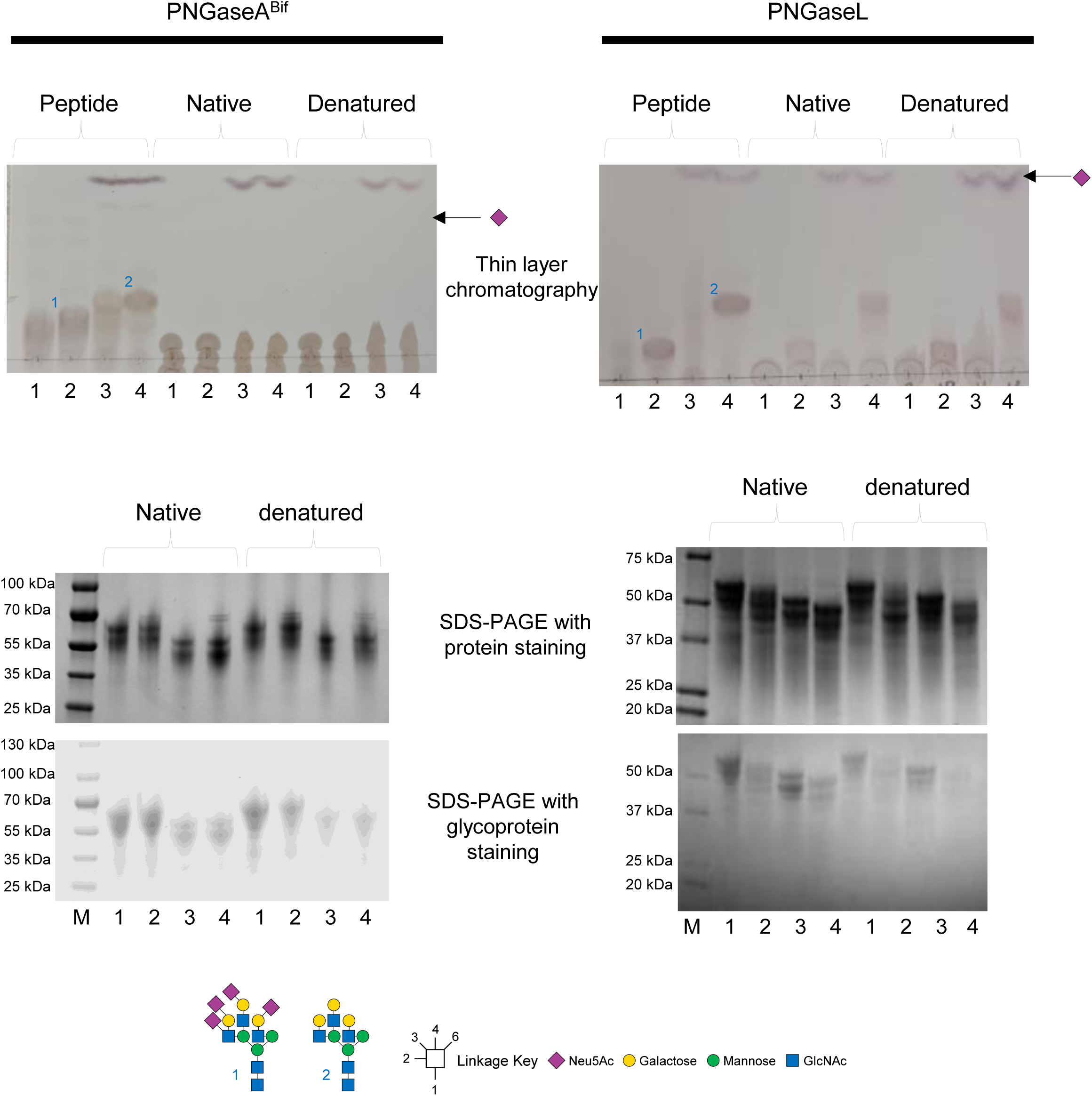
Biochemical Characterisation of PNGaseA^Bif^ against mammalian triantennary complex N-glycans. Bovine fetuin was used to assess the specificity of PNGaseA^Bif^ (left) for mammalian triantennary complex N-glycans and PNGaseL was used as a positive control (right). The peptide was provided to the PNGase enzymes in three forms – native, denatured (boiling), and peptide (*Vibrio coralliilyticus* treated). The impact of removing the α-linked sialic acid from the non-reducing ends of the glycans was also investigated using BT0455 from *Bacteroides thetaiotaomicron*. The N-glycan release was assessed through thin layer chromatography (top) and change in glycoprotein size was monitored through SDS-PAGE with staining for protein (middle) and staining for glycoprotein (bottom). 1: control, 2: PNGase, 3: α-sialidase, 4: PNGase and α-sialidase. M: marker. The *Vibrio coralliilyticus-*derived fragments are too small to resolve by SDS-PAGE.

**Supplementary Figure 6.**
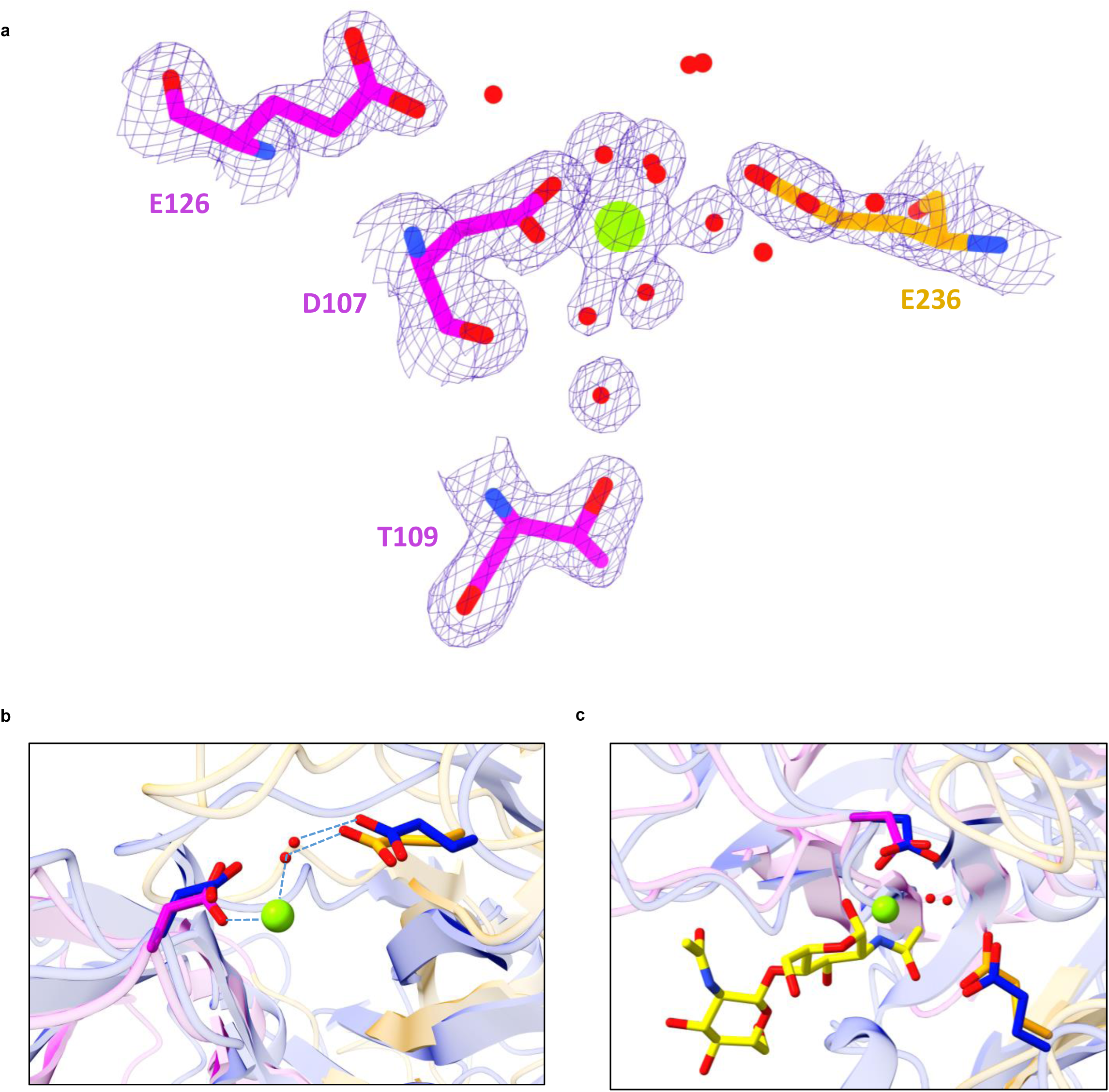
**a)** The 2Fo-Fc electron density map (blue mesh) is shown at 1.5 σ within 1.5 Å distance of E126, D107, E236, T109 and waters involved in the coordination. The electron density strongly supports the octahedral geometry and accurate placement of the water molecules. **b)** Superposition of the conserved Asp and Glu residues shows a coordinating water modelled in the same position in the PNGaseF structure. **c)** Superposition of the chitobiose moity on the PNGaseA^Bif^ structure shows that the magnesium atom in the catalytic site is found in close proximity to where the reducing end of the glycan is expected to be.

**Supplementary Figure 7.**
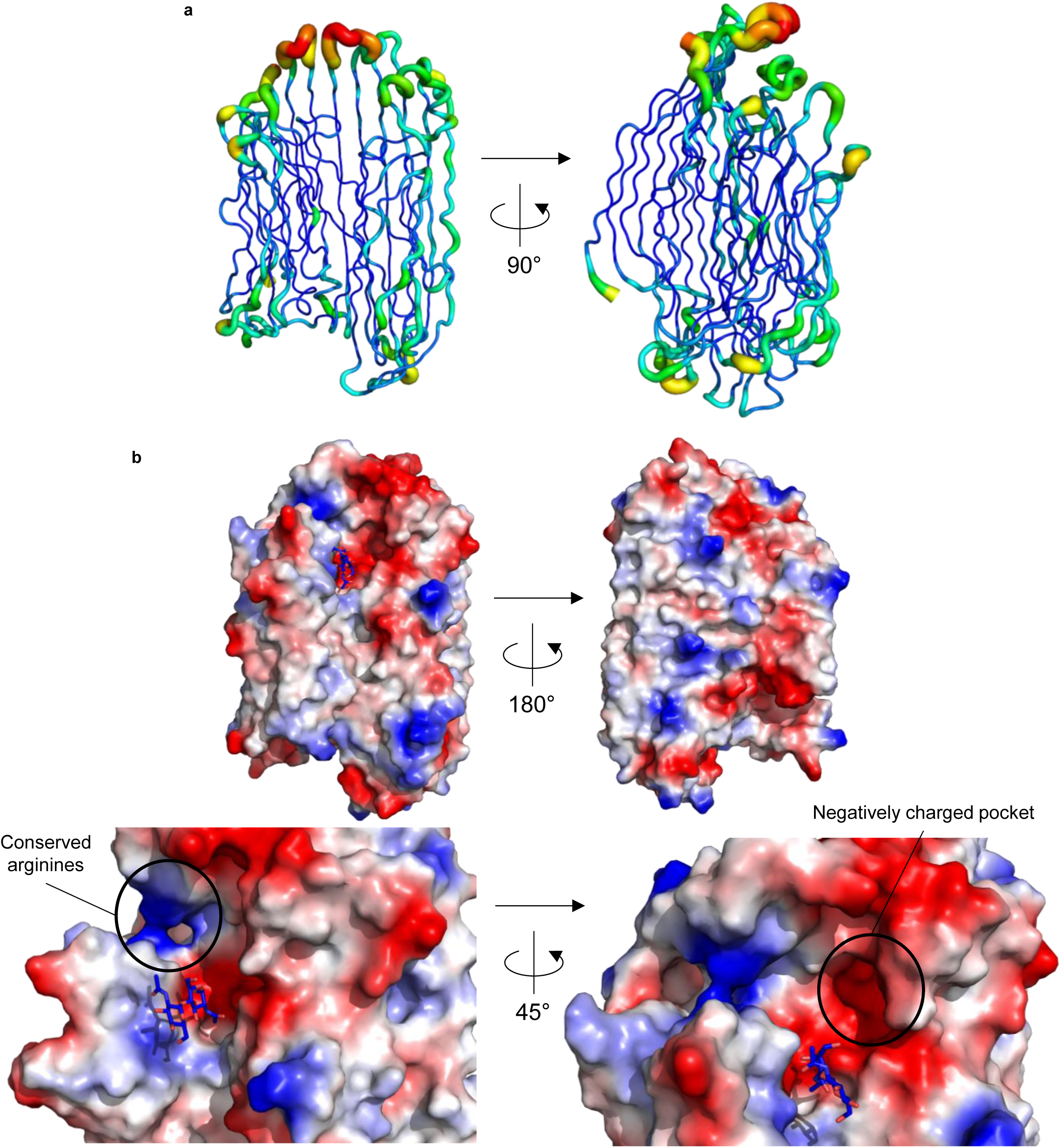
Analysis of the PNGaseA^Bif^ structure. **a)** B-factors of the PNGaseA^Bif^ structure shows that the loops of the cradle nearest the active site have flexibility. b) The electrostatics of the surface of PNGaseA^Bif^ are shown, which highlights the two charged regions in the area that accommodates the peptide component of the substrate.

**Supplementary Figure 8.**
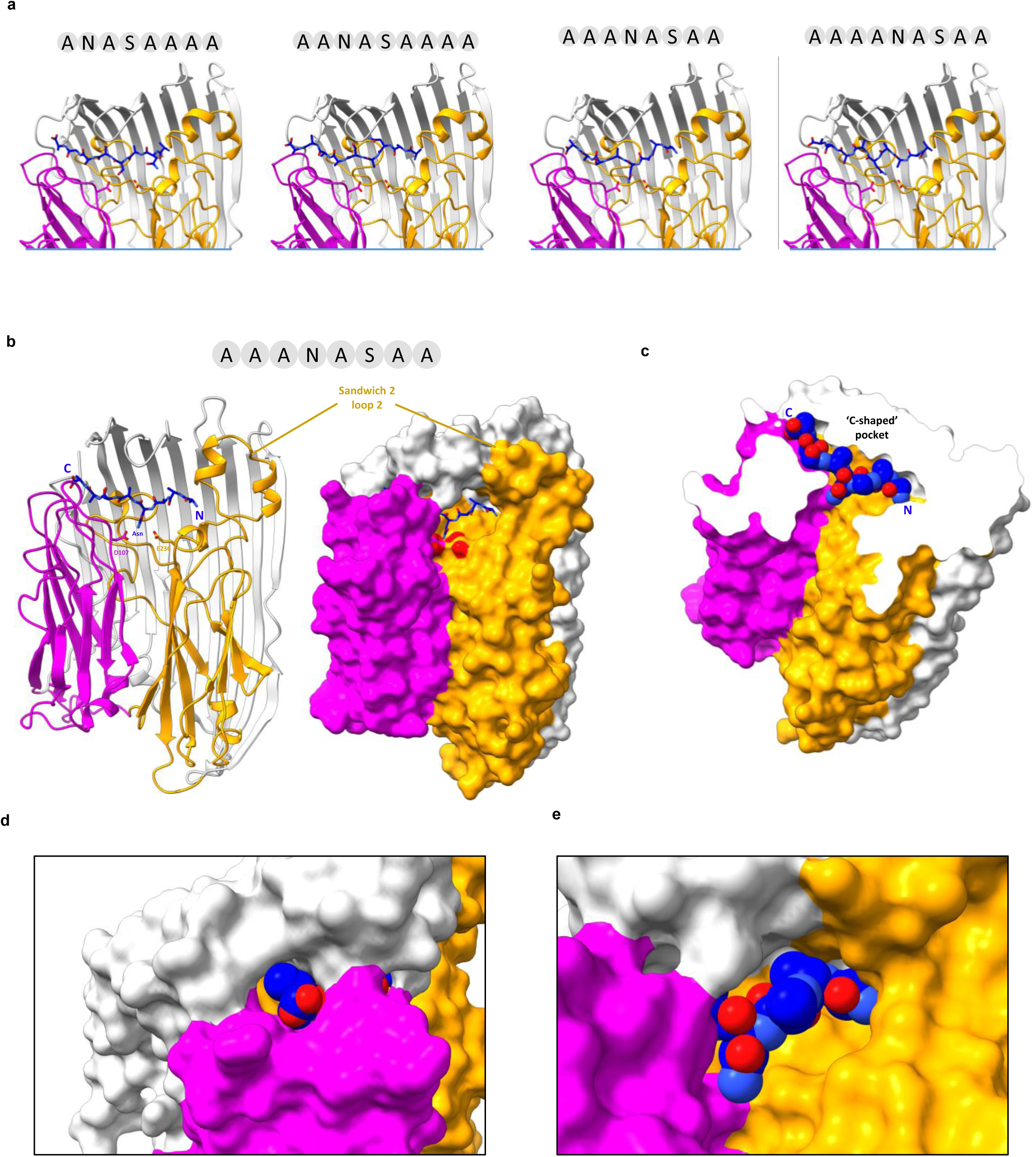
Modelling of peptides into the C-shaped poket. **a)** Four Alphafold 3 models of different peptides (blue and hetero atom colouring) containing the core Asn-Ala-Ser motif. Four predictions are shown with different numbers of Ala residues N- and C-terminal to the NAS motif. Only models with three or four N-terminal residues allow for the N-glycan to reside in proximity to the catalytic residues D107 and E236. **b)** The AAANASAA peptide is shown in the context of the full protein and its encapsulation within the β-sheet cradle. **c)** A clipped model shows the peptide’s accommodation within the ‘C-shaped’ pocket. The pocket would allow for small or hydrophobic N+1, N-2 and N-3 residues. **d)** The C-terminus of the peptide is observed exiting the pocket via the open end of the pocket. **e)** The N-terminus is buried within the closed end of the pocket formed by sandwich 2 loop 2.

**Supplementary Figure 9.**
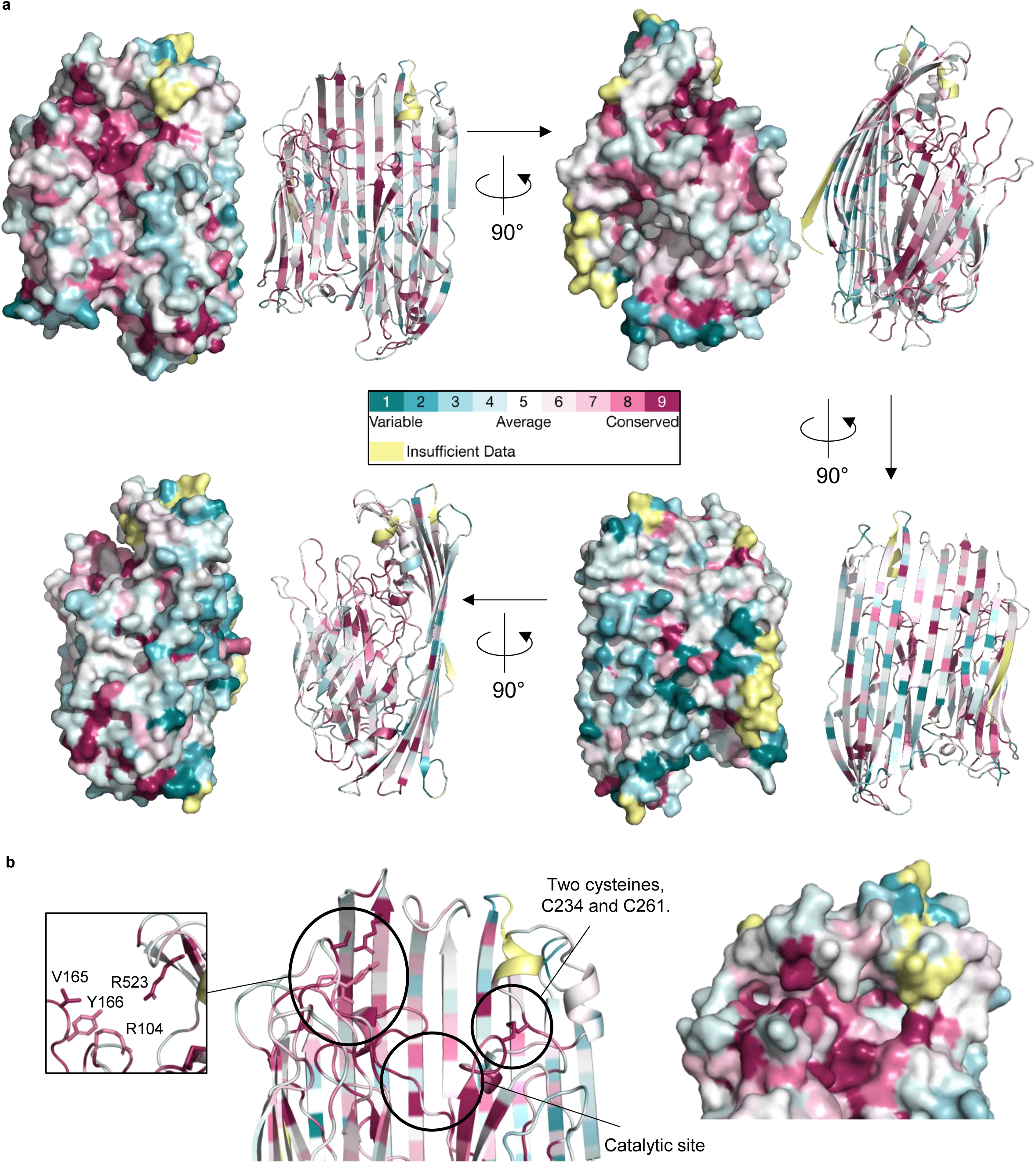
Analysis of the PNGaseA^Bif^ structure. **a)** Consurf was used to analyse the conservation of residues over the PNGaseA^Bif^ structure. The highest conservation of residues is focussed on the active site. b) Consurf analysis of PNGaseA^Bif^ structure that indicates how highly residues are conserved throughout a structure. Three highly conserved regions were identified in the region of the protein where the active site is.

**Supplementary Figure 10.**
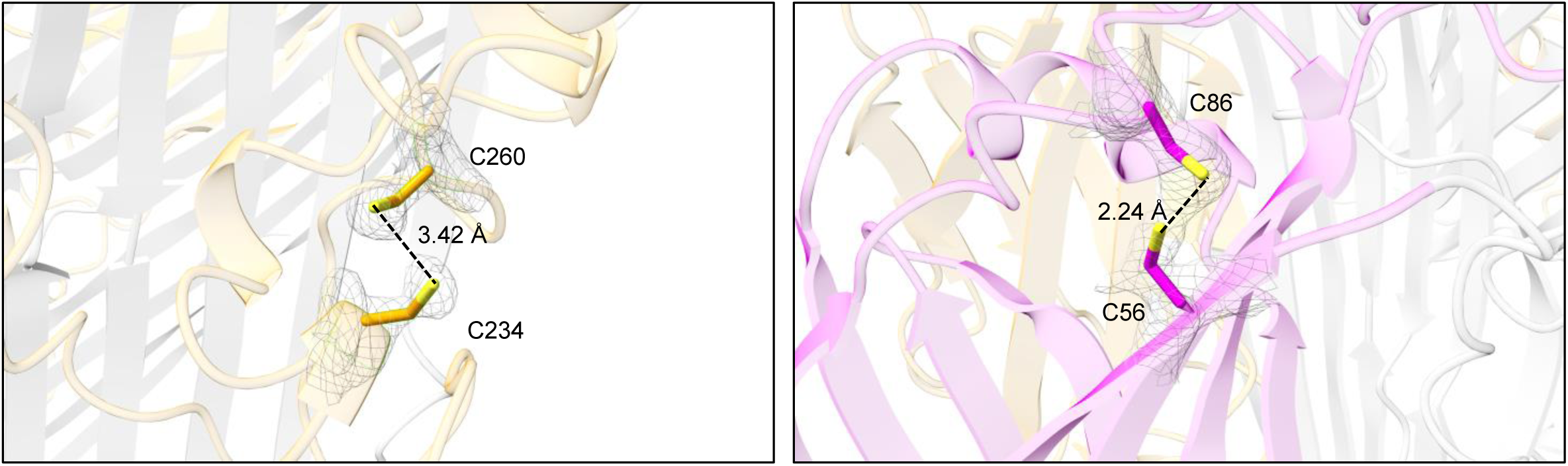
Cysteine/Cystine within PNGase A^bif^. **(a)** The 2Fo-Fc electron density map (mesh) is shown at 2 σ within 2 Å distance of C260 and C234. The density and distance between the sulphur atoms do not support a disulphide bond. **(b)** The 2Fo-Fc electron density map (mesh) is shown at 1.5 σ within 2 Å distance of C56 and C86. The density suggests multiple conformations of the Cys residues, with a 0.58:0.42 ratio between cystine and cysteine forms modelled.

**Supplementary Figure 11.**
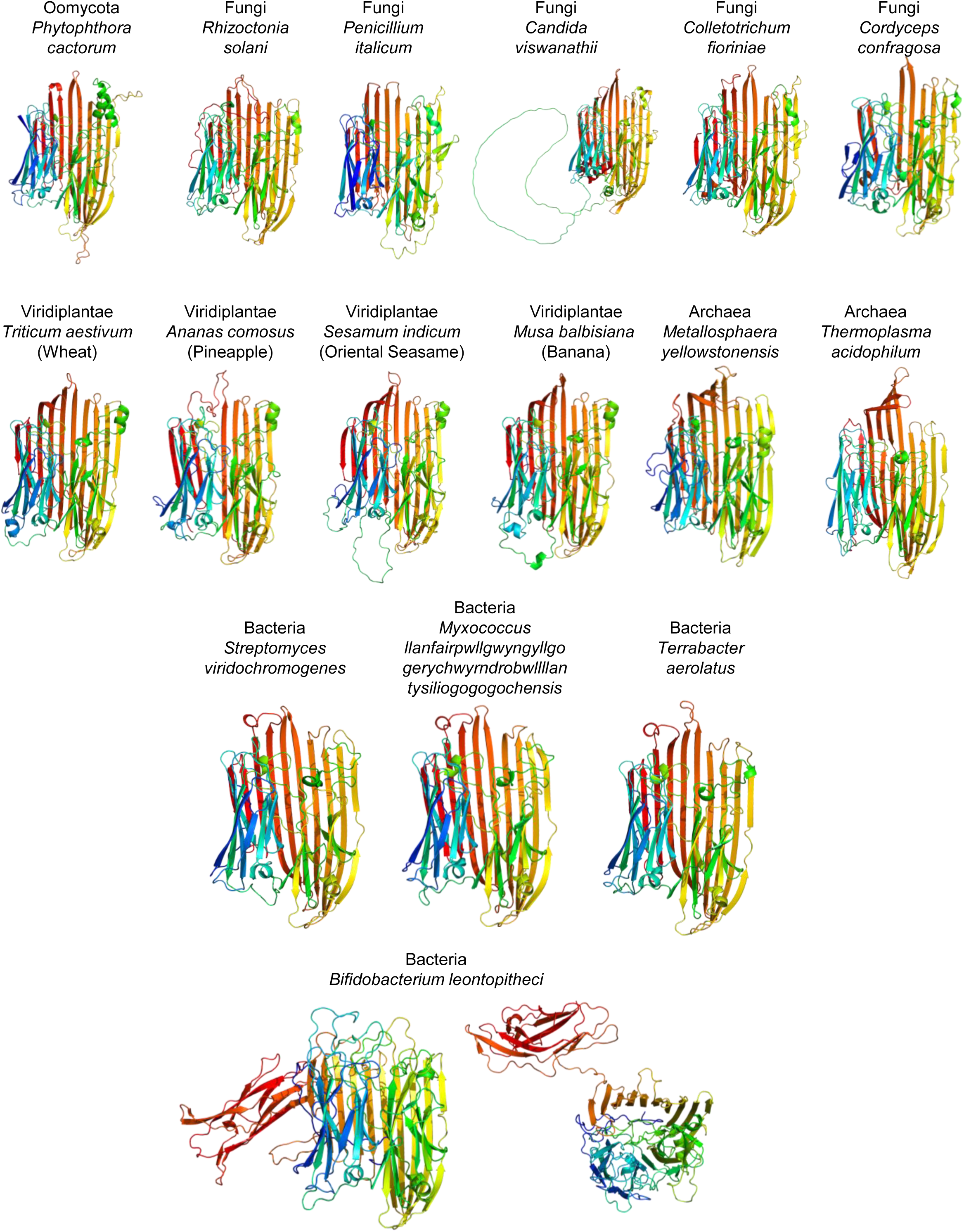
Models of the structures of PNGaseA enzymes from a variety of organisms. The models are displayed as cartoons with the N- and C-termini colours blue to red, respectively. All structures show the canonical two eight-stranded β-sandwich catalytic module and also the cradle, analogous to the PNGaseA^Bif^ structure.

## References

1 Crouch, L. I. N-glycan breakdown by bacterial CAZymes. Essays Biochem 67, 371–383, doi:10.1042/EBC20220256 (2023).

2 Cuskin, F. et al. Human gut Bacteroidetes can utilize yeast mannan through a selfish mechanism. Nature 517, 165–169, doi:10.1038/nature13995 (2015).

3 Briliute, J. et al. Complex N-glycan breakdown by gut Bacteroides involves an extensive enzymatic apparatus encoded by multiple co-regulated genetic loci. Nat Microbiol 4, 1571–1581, doi:10.1038/s41564-019-0466-x (2019).

4 Robb, M. et al. Molecular Characterization of N-glycan Degradation and Transport in Streptococcus pneumoniae and Its Contribution to Virulence. PLoS Pathog 13, e1006090, doi:10.1371/journal.ppat.1006090 (2017).

5 Robb, M. et al. A Second beta-Hexosaminidase Encoded in the Streptococcus pneumoniae Genome Provides an Expanded Biochemical Ability to Degrade Host Glycans. J Biol Chem 290, 30888–30900, doi:10.1074/jbc.M115.688630 (2015).

6 Crouch, L. I. et al. Plant N-glycan breakdown by human gut Bacteroides. Proc Natl Acad Sci U S A 119, e2208168119, doi:10.1073/pnas.2208168119 (2022).

7 Cordeiro, R. L. et al. N-glycan Utilization by Bifidobacterium Gut Symbionts Involves a Specialist beta-Mannosidase. J Mol Biol 431, 732–747, doi:10.1016/j.jmb.2018.12.017 (2019).

8 Stewart, C. J. et al. Temporal development of the gut microbiome in early childhood from the TEDDY study. Nature 562, 583–588, doi:10.1038/s41586-018-0617-x (2018).

9 Garrido, D. et al. Endo-beta-N-acetylglucosaminidases from infant gut-associated bifidobacteria release complex N-glycans from human milk glycoproteins. Mol Cell Proteomics 11, 775–785, doi:10.1074/mcp.M112.018119 (2012).

10 Karav, S. et al. Oligosaccharides Released from Milk Glycoproteins Are Selective Growth Substrates for Infant-Associated Bifidobacteria. Appl Environ Microbiol 82, 3622–3630, doi:10.1128/aem.00547-16 (2016).

11 Cordeiro, R. L. et al. Mechanism of high-mannose N-glycan breakdown and metabolism by Bifidobacterium longum. Nat Chem Biol 19, 218–229, doi:10.1038/s41589-022-01202-4 (2023).

12 Davis, J. C. et al. Identification of Oligosaccharides in Feces of Breast-fed Infants and Their Correlation with the Gut Microbial Community. Mol Cell Proteomics 15, 2987–3002, doi:10.1074/mcp.M116.060665 (2016).

13 Blum, M. et al. InterPro: the protein sequence classification resource in 2025. Nucleic Acids Res 53, D444–D456, doi:10.1093/nar/gkae1082 (2025).

14 UniProt, C. UniProt: the Universal Protein Knowledgebase in 2023. Nucleic Acids Res 51, D523–D531, doi:10.1093/nar/gkac1052 (2023).

15 Chen, I. A. et al. The IMG/M data management and analysis system v.6.0: new tools and advanced capabilities. Nucleic Acids Res 49, D751–d763, doi:10.1093/nar/gkaa939 (2021).

16 van den Belt, M., et al. CAGECAT: The CompArative GEne Cluster Analysis Toolbox for rapid search and visualisation of homologous gene clusters. BMC Bioinformatics 24, 181, doi:10.1186/s12859-023-05311-2 (2023).

17 Abramson, J. et al. Accurate structure prediction of biomolecular interactions with AlphaFold 3. Nature 630, 493–500, doi:10.1038/s41586-024-07487-w (2024).

18 Teufel, F. et al. SignalP 6.0 predicts all five types of signal peptides using protein language models. Nat Biotechnol 40, 1023–1025, doi:10.1038/s41587-021-01156-3 (2022).

19 Sievers, F. et al. Fast, scalable generation of high-quality protein multiple sequence alignments using Clustal Omega. Mol Syst Biol 7, 539, doi:10.1038/msb.2011.75 (2011).

20 Lombard, V., Golaconda Ramulu, H., Drula, E., Coutinho, P. M. & Henrissat, B. The carbohydrate-active enzymes database (CAZy) in 2013. Nucleic Acids Res 42, D490–495, doi:10.1093/nar/gkt1178 (2014).

21 Bakshani, C. R. et al. PNGaseL from Flavobacterium akiainvivens targets a diverse range of N-glycan structures. *BioRxiv*, 10.1101/2025.05.27.656428 (2025).

22 Park, K. H. et al. Structural and biochemical characterization of the broad substrate specificity of Bacteroides thetaiotaomicron commensal sialidase. Biochim Biophys Acta 1834, 1510–1519, doi:10.1016/j.bbapap.2013.04.028 (2013).

23 Zhang, Z., Xie, J., Zhang, F. & Linhardt, R. J. Thin-layer chromatography for the analysis of glycosaminoglycan oligosaccharides. Anal Biochem 371, 118–120, doi:10.1016/j.ab.2007.07.003 (2007).

24 Mirdita, M. et al. ColabFold: making protein folding accessible to all. Nat Methods 19, 679–682, doi:10.1038/s41592-022-01488-1 (2022).

25 Adams, P. D. et al. PHENIX: a comprehensive Python-based system for macromolecular structure solution. Acta Crystallogr D Biol Crystallogr 66, 213–221, doi:10.1107/s0907444909052925 (2010).

26 Emsley, P., Lohkamp, B., Scott, W. G. & Cowtan, K. Features and development of Coot. Acta Crystallogr D Biol Crystallogr 66, 486–501, doi:10.1107/s0907444910007493 (2010).

27 The PyMOL Molecular Graphics System, Version 2.0 Schrödinger, LLC.

28 Ben Chorin, A., et al. ConSurf-DB: An accessible repository for the evolutionary conservation patterns of the majority of PDB proteins. Protein Sci 29, 258–267, doi:10.1002/pro.3779 (2020).

29 Duranti, S. et al. Characterization of the phylogenetic diversity of two novel species belonging to the genus Bifidobacterium: Bifidobacterium cebidarum sp. nov. and Bifidobacterium leontopitheci sp. nov. Int J Syst Evol Microbiol 70, 2288–2297, doi:10.1099/ijsem.0.004032 (2020).

30. Arzamasov, A. A., et al. Integrative genomic reconstruction reveals heterogeneity in carbohydrate utilization across human gut bifidobacteria. bioRxiv, doi:10.1101/2024.07.06.602360 (2025).

31 Letunic, I. & Bork, P. 20 years of the SMART protein domain annotation resource. Nucleic Acids Res 46, D493–d496, doi:10.1093/nar/gkx922 (2018).

32 Altmann, F., Schweiszer, S. & Weber, C. Kinetic comparison of peptide: N-glycosidases F and A reveals several differences in substrate specificity. Glycoconj J 12, 84–93, doi:10.1007/BF00731873 (1995).

33 Mass, S. et al. The coral pathogen Vibrio coralliilyticus uses a T6SS to secrete a group of novel anti-eukaryotic effectors that contribute to virulence. PLoS Biol 22, e3002734, doi:10.1371/journal.pbio.3002734 (2024).

34 Sun, G. et al. Identification and characterization of a novel prokaryotic peptide: N-glycosidase from Elizabethkingia meningoseptica. J Biol Chem 290, 7452–7462, doi:10.1074/jbc.M114.605493 (2015).

35 Kuhn, P. et al. Active site and oligosaccharide recognition residues of peptide-N4-(N-acetyl-beta-D-glucosaminyl)asparagine amidase F. J Biol Chem 270, 29493–29497, doi:10.1074/jbc.270.49.29493 (1995).

36 Bakshani, C. R. et al. Carbohydrate-active enzymes from Akkermansia muciniphila break down mucin O-glycans to completion. Nat Microbiol 10, 585–598, doi:10.1038/s41564-024-01911-7 (2025).

37 Gotoh, A. et al. Novel substrate specificities of two lacto-N-biosidases towards beta-linked galacto-N-biose-containing oligosaccharides of globo H, Gb5, and GA1. Carbohydr Res 408, 18–24, doi:10.1016/j.carres.2015.03.005 (2015).

38 Miwa, M. et al. Cooperation of beta-galactosidase and beta-N-acetylhexosaminidase from bifidobacteria in assimilation of human milk oligosaccharides with type 2 structure. Glycobiology 20, 1402–1409, doi:10.1093/glycob/cwq101 (2010).

39 Takada, H., Katoh, T., Sakanaka, M., Odamaki, T. & Katayama, T. GH20 and GH84 beta-N-acetylglucosaminidases with different linkage specificities underpin mucin O-glycan breakdown capability of Bifidobacterium bifidum. J Biol Chem 299, 104781, doi:10.1016/j.jbc.2023.104781 (2023).

40 Katoh, T. et al. Identification and characterization of a sulfoglycosidase from Bifidobacterium bifidum implicated in mucin glycan utilization. Biosci Biotechnol Biochem 81, 2018–2027, doi:10.1080/09168451.2017.1361810 (2017).

41 Wakinaka, T. et al. Bifidobacterial alpha-galactosidase with unique carbohydrate-binding module specifically acts on blood group B antigen. Glycobiology 23, 232–240, doi:10.1093/glycob/cws142 (2013).

42 Kiyohara, M. et al. An exo-alpha-sialidase from bifidobacteria involved in the degradation of sialyloligosaccharides in human milk and intestinal glycoconjugates. Glycobiology 21, 437–447, doi:10.1093/glycob/cwq175 (2011).

43 Sela, D. A. et al. An infant-associated bacterial commensal utilizes breast milk sialyloligosaccharides. J Biol Chem 286, 11909–11918, doi:10.1074/jbc.M110.193359 (2011).

44 Nishiyama, K. et al. Two extracellular sialidases from Bifidobacterium bifidum promote the degradation of sialyl-oligosaccharides and support the growth of Bifidobacterium breve. Anaerobe 52, 22–28, doi:10.1016/j.anaerobe.2018.05.007 (2018).

45 Sakurama, H. et al. Differences in the substrate specificities and active-site structures of two alpha-L-fucosidases (glycoside hydrolase family 29) from Bacteroides thetaiotaomicron. Biosci Biotechnol Biochem 76, 1022–1024, doi:10.1271/bbb.111004 (2012).

46 Katayama, T. et al. Molecular cloning and characterization of Bifidobacterium bifidum 1,2-alpha-L-fucosidase (AfcA), a novel inverting glycosidase (glycoside hydrolase family 95). J Bacteriol 186, 4885–4893, doi:10.1128/jb.186.15.4885-4893.2004 (2004).

47 Ambrogi, V. et al. Characterization of GH2 and GH42 beta-galactosidases derived from bifidobacterial infant isolates. AMB Express 9, 9, doi:10.1186/s13568-019-0735-3 (2019).

48 Sakurama, H. et al. Lacto-N-biosidase encoded by a novel gene of Bifidobacterium longum subspecies longum shows unique substrate specificity and requires a designated chaperone for its active expression. J Biol Chem 288, 25194–25206, doi:10.1074/jbc.M113.484733 (2013).

49 Shimada, Y. et al. alpha-N-Acetylglucosaminidase from Bifidobacterium bifidum specifically hydrolyzes alpha-linked N-acetylglucosamine at nonreducing terminus of O-glycan on gastric mucin. Appl Microbiol Biotechnol 99, 3941–3948, doi:10.1007/s00253-014-6201-x (2015).

50 Barratt, M. J. et al. Bifidobacterium infantis treatment promotes weight gain in Bangladeshi infants with severe acute malnutrition. Sci Transl Med 14, eabk1107, doi:10.1126/scitranslmed.abk1107 (2022).

